# From Latent Manifolds to Functional Probes: An Interpretable, Kinome-Scale Generative Machine Learning Framework for Family-Targeted Kinase Inhibitor Design

**DOI:** 10.64898/2026.01.06.698033

**Authors:** Ryan Kassab, Keerthi Krishnan, Gennady Verkhivker

## Abstract

The design of selective kinase inhibitors remains a formidable challenge due to the high structural conservation of the ATP-binding site across the kinome. While modern generative AI has enabled rapid exploration of chemical space, many advanced models operate as black boxes, obscuring the chemical rationale behind design choices and limiting interpretability. To explore these bottlenecks, we present a modular, generative framework for *de novo* design of SRC kinase inhibitors, integrating ChemVAE-based latent space modeling, a chemically interpretable Kinase Inhibition Likelihood (KIL) scoring function, Bayesian optimization, and cluster-guided local neighborhood sampling. The results demonstrate that kinase inhibitors spontaneously organize into a coherent, low-dimensional manifold in latent space, with SRC acting as a structural “hub” that enables rational scaffold transformation. Our local neighborhood sampling-based approach successfully converts inhibitors from other kinase families (notably LCK) into novel SRC-like chemotypes, with LCK-derived molecules accounting for ∼40% of high-similarity outputs. Critically, we expose a fundamental representation gap: despite aromatic ring count being a top KIL feature, SMILES-based generation systematically fails to access multi-ring pharmacophores characteristic of clinical kinase inhibitors. This limitation cannot be overcome by scoring refinement alone, demanding topology-aware representations. Our framework also demonstrates that unbiased exploration paired with cluster-guided sampling outperforms active-biased optimization, which traps search in narrow local optima. By exposing representational gaps and showcasing scaffold-aware navigation of latent space, this study argues for hybrid systems that combine the diagnostic transparency of interpretable machine learning frameworks with the generative power of modern architectures.

## 1. Introduction

Discovery of small-molecule inhibitors—especially against high-value but structurally complex targets such as kinases, GPCRs, and protein–protein interfaces—unfolded through a painstaking cycle of chemical synthesis, high-throughput screening, and iterative structure–activity relationship (SAR) analysis. This process, constrained by experimental throughput and human intuition, often took years to yield a single clinical candidate. The past decade, however, has witnessed a profound transformation: the integration of artificial intelligence (AI) and machine learning (ML) into drug discovery has enabled the *de novo*, property-driven generation of drug-like molecules with unprecedented speed, scale, and chemical novelty [1–8]. Many deep learning approaches have been put forward employing various neural network architectures, molecular representations and analysis metrics for targeted compound design, and their applications [9–16]. This paradigm shift in the drug discovery evolved from syntax-aware sequence modeling toward structure- and function-aware molecular design. The first wave of this transformation emerged when several studies began treating molecules as textual sequences using the Simplified Molecular Input Line Entry System (SMILES) and applying natural language processing (NLP) techniques to chemical space. Deep neural network (DNN) models, most notably variational autoencoder (VAE) [9] and generative adversarial networks (GAN) [17] have been particularly fruitful in molecular design of novel chemical probes [17–28]. Among the earliest efforts was sequence data generation (SeqGAN) approach [18], and Objective-Reinforced Generative Adversarial Networks (ORGAN) [19] which coupled a recurrent neural network (RNN) generator with a discriminator trained not just to assess chemical validity but to maximize user-defined molecular properties—a pioneering step toward goal-directed generation. LatentGAN combined an autoencoder and a generative adversarial neural network for de novo molecular design [20]. DruGAN approach combined GAN and VAE by training an adversarial autoencoder to efficiently sample molecules from the latent space [23]. Soon after, MolGAN [29] adapted GANs to molecular design, generating adjacent and feature matrices to represent molecular graphs directly. CycleGAN provided unpaired Image-to-Image translation using Cycle-Consistent Adversarial Networks [30]. MolCycleGAN, which extended the CycleGAN framework, can learn transformation rules from the sets of compounds with desired and undesired values of the considered property [31]. The methodological progress in GAN applications to molecular discovery has been catalyzed by the development of several comprehensive benchmarking sets and cheminformatics infrastructure [32–36]. Despite its conceptual elegance, MolGAN and related GAN approaches suffered from severe mode collapse and produced valid molecules less than 30% of the time, highlighting the fragility of adversarial training in discrete spaces. A more robust alternative arrived with ChemVAE [9], a variational autoencoder that encoded SMILES strings into a smooth, continuous 196-dimensional latent space while simultaneously predicting key drug-likeness metrics—quantitative estimate of drug-likeness (QED) [37], synthetic accessibility score (SAS) [38], and logP [39].

Subsequent reward-guided strategies such as REINVENT approach [20,40,41] proposed a reinforcement learning (RL) framework that used policy gradients to fine-tune and RNN generator toward a customizable reward function. This reward could combine multiple objectives—such as predicted binding affinity, QED, and Tanimoto similarity to a reference scaffold—effectively turning the generative model into a programmable design engine. GENTRL approach compresses the space of small molecule structures onto a distribution that parameterizes the latent space in a high-dimensional lattice following by exploration and optimization of the latent space by reinforcement learning to discover novel kinase inhibitors [42].

Concurrently, transformer architectures began to reshape the landscape of chemical AI. Attention-based generative models for de novo molecular design offered new architectures that enabled a more accurate sampling from the latent space and exploration of novel chemistry space not present in the training data [43], thus optimizing the tradeoffs between model exploration and structure of the latent memory. Efficient multi-objective molecular design approaches combine *in silico* prediction of molecular properties defined desirability ranges and substructure constraints with particle swarm optimization for optimal navigation in a continuous latent space [44–46]. A highly efficient and generic query-based molecule optimization framework QMO facilitates molecule optimization by decoupling molecule representation learning and guided search method based on zeroth-order optimization in the molecular property landscape [47]. The performances of DL-based, VAE, GAN and RNN models were evaluated in goal-directed (rediscovery, optimization and scaffold hopping of active compounds) and target-specific (generation of novel compounds for a given target) tasks [48]. Simultaneously, SMILES-BERT [49] and Chemformer [50,51] applied transformer architectures to molecular sequences, leveraging self-supervised pretraining on billions of compounds to improve generation quality and transfer learning. These approaches offered greater controllability and higher validity, but remained constrained by the sequential nature of SMILES, which struggles to represent cyclic and stereochemical complexity. MolDQN bypassed SMILES by applying deep Q-learning to discrete molecular graph actions, optimizing molecular properties through a Markov decision process [52]. Despite these advances, this phase remained two-dimensional: rewards were often computed using surrogate predictors—such as Random Forests trained on RDKit descriptors rather than direct protein–ligand interactions, yielding molecules that were chemically plausible but often pharmacologically inert.

Some generative models aiming at three-dimensional (3D) molecule generation have also been proposed, gaining attention for their unique advantages and potential to explicitly design drug-like molecules in a target-conditioning manner [53]. A novel molecular deep generative model adopts a recurrent neural network architecture coupled with a ligand-protein interaction fingerprint as constraints [54]. DeepLigBuilder, a deep learning-based method for *de novo* drug design combined Ligand Neural Network (L-Net) graph generative model for design of chemically and conformationally valid 3D molecules with Monte Carlo tree search to optimize structure-based *de novo* drug design parameters such as high predicted affinity, and similar binding features to those of known inhibitors [55]. A comprehensive review of 3D molecular generative models reported current techniques for the molecular structure generation and categorized them into three types, depending on featurization of 3D molecular structures: cubic grid-based, Euclidean distance matrix(EDM)-based, and Cartesian coordinate-based, where each type of featurization requires distinct generative architectures and optimization strategies [56].

Th advent of graph representation learning, a class of ML methods that natively operate on graph-structured data, has enabled neural networks to learn directly from molecular topology. Over the past seven years, graph neural networks (GNNs) have redefined the very architecture of *de novo* molecular design, virtual screening, and protein–ligand interaction modeling. The adoption of GNNs in drug discovery began with their ability to natively represent molecules as graphs and learn structure–property relationships directly from topology. Gilmer et al. [57] laid out the conceptual foundation of GNNs with the introduction of Neural Message Passing (MPNN), a unifying framework that cast molecular property prediction as an iterative process of information exchange between atoms and bonds. This insight catalyzed a wave of chemistry-specific GNN architectures. Kearnes et al. demonstrated that Graph Convolutional Networks (GCNs) could predict ADMET properties with high accuracy by aggregating neighborhood features in molecular graphs [58]. Soon after, Veličković et al. introduced Graph Attention Networks (GATs), which learned to weight the importance of neighboring atoms dynamically, capturing subtle electronic effects critical for reactivity and binding [59]. A pivotal advance came with Directed Message Passing Neural Networks (D-MPNN) approach which explicitly modeled bond directionality and chirality—features essential for drug-likeness and target specificity [60] D-MPNN achieved state-of-the-art performance across quantum chemical (QM9) and bioactivity (MUV, Tox21) benchmarks and became the core of the open-source Chemprop framework, now widely adopted in both academia and industry for interpretable molecular property prediction [60].JT-VAE approach widely adopted by 2021, decomposed molecules into hierarchical junction trees of rings and chains, enabling near-perfect validity (>99%) and precise control over scaffold modification [61]. Junction tree variational autoencoder (JT-VAE) generates molecular graphs in two phases, by first generating a tree-structured scaffold over chemical substructures and then combining them into a molecule with a graph message passing network [61]. Similarly, GraphAF used autoregressive normalizing flows to build molecular graphs atom-by-atom with high fidelity, ensuring that valency and chirality were respected [62]. More recently, GFlowNets [63,64] introduced a probabilistic framework for sampling molecules proportional to a reward function (e.g., binding affinity), mitigating the mode collapse and low diversity that plagued earlier generative approaches.

The limitations of static, 2D graph representations particularly for tasks required 3D conformational awareness, such as protein–ligand docking and allosteric modulation. This spurred the rise of geometric deep learning, where models respect the rotational and translational symmetries of physical space. SchNet, introduced by Schütt et al. [65,66] pioneered the use of continuous-filter convolutions operating directly on atomic coordinates, enabling accurate prediction of molecular energies and interatomic forces with quantum-mechanical fidelity. This was significantly refined in DimeNet and DimeNet++ [67,68] which incorporated directional message passing using interatomic angles, dramatically improving the modeling of torsional strain, steric clashes, and binding pocket complementarity. The culmination of this trend arrived with SE(3)-equivariant GNNs, exemplified by EquiBind which predicted protein–ligand binding poses in seconds—without traditional docking—by learning geometric constraints directly from structural data [69]. EquiBind achieved near-experimental accuracy on the PDBBind benchmark, effectively replacing physics-based scoring in early-stage screening [69].

Critically, graph representation learning also enabled direct modeling of protein–ligand complexes as heterogeneous graphs, where protein residues and ligand atoms form distinct node types connected by cross-edges. TANKBind approach segments the whole protein into functional blocks and predict their interactions with the ligand, creating a protein-ligand interaction energy landscape using a novel trigonometry-aware architecture. In the second stage, TANKBind prioritizes the crystal structures by contrastively ensuring a weaker binding affinity for non-native interactions [70]. Self-supervised, pretrainable geometric GNNs can learn rich representations of molecules and proteins from unlabeled structural data and represent a new class of models designed for molecular property prediction that leverage 3D molecular structure information during pre-training to improve performance on downstream tasks [71–73]. Graph Multi-View Pre-training (GraphMVP) framework addresses limited 3D molecular data by using self-supervised learning with contrastive learning to enforce consistency between 2D and 3D molecular representations, enhancing performance in property prediction tasks [71].

The current frontier integrates generative modeling and massive scale pretraining into end-to-end systems capable of co-designing proteins and ligands from first principles. Diffusion models have emerged as one of the dominant generative frameworks. GeoDiff is the first SE(3)-equivariant diffusion model that operates directly on atomic coordinates and learns to reverse a diffusion process that gradually adds noise to a molecule’s 3D structure [74]. This approach enabled structure-aware ligand generation for docking and binding prediction and laid the foundation for protein-conditioned diffusion models. By expanding this work, Tang and his group introduced a pretrainable, SE(3)-equivariant geometric GNN specifically designed for antibody affinity maturation [75]. This work bridged geometric deep learning and self-supervised pretraining with high-throughput experimental biology to create a predictive, generative, and actionable platform for antibody engineering. These studies underscored a broader trend: the shift from sequence-based to structure-based AI in therapeutic discovery, with geometric GNNs at the forefront. TorsionDiff [76] operates in torsion angle space to produce realistic side-chain rotamers. DiffDock predicts binding poses with near-experimental accuracy, effectively replacing classical docking pipelines [77]. DiffDock-L, a latest version of DiffDock provides a significant improvement in performance and generalization capacity [78]. DiffLinker is a new Equivariant 3D-conditional Diffusion Model for Molecular Linker Design that places missing atoms in between and designs a molecule incorporating all the initial fragments [79].

When conditioned on protein structure, diffusion-based models achieve remarkable biological specificity. RFdiffusion approach developed by the Baker Lab uses a protein backbone diffusion model [80] and when paired with the sequence design tool ProteinMPNN [81] enables *de novo* creation of protein binders, allosteric pockets, and small-molecule scaffolds. Complementing these are foundation models trained on multimodal biological data. ESM3 integrates sequences, 3D structures, and functional annotations into a single architecture capable of zero-shot ligand generation via in-context learning [82]. Chroma, a generative model for proteins and protein complexes, can directly sample novel protein structures and sequences, and that can be conditioned to steer the generative process towards desired properties and functions [83]. This evolution has been accelerated by open-source ecosystems and standardized benchmarks. PyTorch Geometric (PyG) [84] and Deep Graph Library (DGL) [85] provide modular, scalable implementations of GNN layers. The Therapeutics Data Commons (TDC) [86] offers a unified benchmark with 66 therapeutic tasks, including kinase inhibitor design, antibody escape, and molecular generation.

In current generative AI approaches, scoring is embedded within the generative process itself, while multi-objective trade-offs between potency and selectivity are handled implicitly through conditional diffusion or multi-task pretraining. However, while modern AI tools offer unprecedented generative power they often operate as black boxes, making it difficult to diagnose failure modes, ensure pharmacophoric fidelity or guide iterative refinement. Hybrid approaches, combining the predictive power of GNNs with chemically grounded, interpretable features metric, may help to address this important gap. In the development of kinase inhibitory therapeutics, generating novel selective probes to interrogate specific protein kinases remains a considerable challenge and ML-enabled targeted transformations and chemical morphing between kinase inhibitors from different families can provide a valuable resource for new indications of existing kinase molecules.

In this study, we present a modular, interpretable ML-based platform for the *de novo* design of SRC kinase inhibitors. Our generative pipeline employs a hybrid AI framework that integrates deep variational autoencoding, interpretable ML–based scoring, and probabilistic optimization to enable targeted exploration of kinase inhibitor chemical space. The presented strategy decomposes the design process into several interconnected stages : (a) deep generative backbone for latent space representation, (b) ML scorer for target-specific guidance, (c) probabilistic optimization engine (Bayesian Optimization) for search, (d) clustering-and-local neighborhood sampling layer for scaffold transformation. This work represents a significant conceptual and methodological extension of our earlier study [87]. While our earlier study demonstrated initial scaffold transformation via latent space local neighborhood sampling, the present work delivers a more comprehensive, multi-strategy generative pipeline with several critical advances. Most notably, we now expand the data sets of random molecules and kinase inhibitors, benchmark Bayesian Optimization as a complementary global search strategy revealing its strengths (efficient drug-likeness tuning) and fundamental limitations (systematic failure to recover multi-ring pharmacophores). The current study offers a mechanistic interpretation of latent space organization, identifying SRC as a structural “hub” and LCK as a uniquely “plastic” scaffold for transformation.

## 2. Materials and Methods

### 2.1. Data Sets of Protein Kinase Inhibitors and Small Molecules

To construct a robust and representative foundation for generative kinase inhibitor design, we assembled a large-scale, multi-source dataset that reflects the current state of kinase-targeted chemical space. Numerous large databases are available that contain molecules in a variety of representations including SMILES, 2D, and 3D. For this study, we explored the databases of generic small molecules and drug-like inhibitors primarily ChEMBL [88], DrugBank [89,90], BindingDB [91], BindingMoad [92], ChEBI [93], ZINC, a free database of commercially available compounds that contains over 230 million purchasable compounds in ready-to-dock, 3D formats [94–96]. Our inhibitor collection integrates high confidence bioactive compounds from ChEMBL v32, DrugBank v5.1 [90], PDBbind v2023 [91] and ZINC20 [96]. To provide a meaningful contrast to kinase-biased chemistry, we sampled drug-like matter from two ultra-large enumerative databases: GDB-17 Lead-Like Set: ∼11 million molecules filtered for lead-like properties (MW ≤ 450, logP ≤ 4, ≤4 HBD/HBA) [97,98], FDB-17 subset ∼10 million fragment-like compounds derived from GDB-17 using synthetic accessibility and complexity filters [99]. From these, we selected ∼ 220,000 diverse molecules satisfying Lipinski’s Rule of Five (MW < 700, logP ∈ [–4,6], ≤6 rotatable bonds, ≤12 HBD/HBA) and restricted to biologically relevant atoms (C, N, O, F, S, P, Cl, Br, I). This “random” background set ensures the model learns to distinguish kinase-specific pharmacophores from generic drug-like space.

For generative kinase inhibitor design, we assembled a comprehensive dataset of protein kinase inhibitors (PKIs). In 2023, Bajorath reported a total of 155,579 qualifying unique human PKIs [100]. Our curation strategy is informed by recent systematic analyses of the kinome-wide inhibitor landscape, including the landmark 2025 review by Koch, Kullmann, and Bajorath [101] which reports that over 206,000 protein kinase inhibitors have been disclosed spanning orthosteric, allosteric, and covalent mechanisms across the human kinome. For datasets of PKIs, we used ∼60,000 available high-confidence PKIs. The expanded set covered the expanded set of kinase families totaling 37 distinct kinase families across the human kinome, including: SRC (SRC, LCK, FYN, YES), ABL (ABL1, ABL2), EGFR (EGFR, ERBB2/HER2, ERBB4), PDGFR (PDGFRα, PDGFRβ, KIT, CSF1R, FLT3), FGFR (FGFR1–4), INSR (INSR, IGF1R), TRK (NTRK1/2/3), ROS (ROS1, DDR1, DDR2), MET (MET, RON), RAF (ARAF, BRAF, CRAF), MLK (MAP3K9, MAP3K10, MAP3K11), LRRK (LRRK1, LRRK2), STKR (ALK, LTK, ROS, RYK), TLK (TLK1, TLK2), RIPK (RIPK1–4), WNK (WNK1–4), CLK (CLK1–4), STE20 (PAK1–7, MAP4K1–7) STE11 (MAP3K1–13), STE7 (MAP2K1–7), CAMK (CAMK1–4, DAPK1–3), DAPK, PHK (PHKG1/2), MLCK (MYLK), DCAMKL (DCAMKL1–3), MELK, BRSK, PKA (PRKACA/B/C), PKG (PRKG1/2), PKC (PRKCA–Z), AKT (AKT1–3), RSK (RPS6KA1–6), SGK (SGK1–3) CDK (CDK1–20), MAPK (MAPK1/3/8/9/11/14/p38α–δ),GSK3 (GSK3A/B).

In the earlier study [87[ we used the data set of competitive and allosteric protein kinase inhibitors confirmed by X-ray crystallography that contained a total of 2,899 unique inhibitors including 136 allosteric and 2763 orthosteric compounds with a total of 231 protein kinases [102–104]. In the current study, we included the latest data from the KLIFS website (accessed April 2025) that reported 4,179 unique ligands confirmed by X-ray across 6,738 structures for 326 kinases [105].

We also expanded the list of allosteric kinase ligands based on recent systematic analysis of X-ray structures that identified a total of 262 allosteric PK ligands [106]. For focused generative experiments on SRC, we extracted 3,477 high-confidence SRC inhibitors (IC ≤ 100 nM) and 1,883 ABL1 inhibitors as reference scaffolds. All molecules were standardized using RDKit [107,108] with salts removed, tautomers normalized, and stereochemistry preserved. All molecules including both kinase inhibitors and background compounds were converted to canonical SMILES and encoded into a 196-dimensional continuous latent space using the ChemVAE architecture [9]. ChemVAE converts discrete representations of molecules to and from a multidimensional continuous representation, enabling generation of new molecules for efficient exploration and optimization via open ended chemical spaces, enabling Bayesian optimization in latent space and allowing to navigate toward regions enriched for desired properties.

### 2.2. Guided Remodeling of Latent Neighborhoods via Cluster-Directed Sampling

To enable scaffold-aware transformation of kinase inhibitors across families, we developed a guided latent space remodeling strategy that leverages the intrinsic structural organization of the ChemVAE embedding. Rather than applying global or random modifications, our approach performs targeted local neighborhood sampling—a process that shifts molecular representations toward chemically coherent regions of latent space while preserving scaffold integrity. We began by applying K-means clustering to the 196-dimensional ChemVAE latent space to identify functionally homogeneous neighborhoods. This unsupervised step avoids manual labeling and allows molecular embeddings to self-organize into groups based solely on structural and physicochemical similarity. We evaluated cluster configurations ranging from 2 to 5 partitions and found that a 3-cluster split yielded the highest diversity and validity of generated molecules, as well as the clearest separation of scaffold motifs (e.g., fused heterocycles vs. linear aromatics). This configuration was selected for all subsequent remodeling experiments. Within each cluster, we performed centroid-directed sampling: for every molecule with latent representation: : for every molecule with latent representation x, we computed its displacement toward the cluster centroid c using a controlled interpolation:

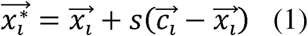

where the scaling factor governs the degree of remodeling. Given that the lower bound of *s* = 0 corresponds to the original encoding of a given molecule, while *s* = 1 provides us with the centroid of the cluster, this parameter was initially set to be a threshold of 0.5. By performing local sampling steps and evaluating kinase inhibition likelihood probabilities, we found that with the scaling factor *s* < 0.5 the yield of valid molecules decreased, while a scaling factor *s* = 0.8 remodels the molecule gradually towards the centroid of the cluster yielding valid molecules without losing information of the molecular attributes. To encourage diversity without destabilizing the latent geometry, we introduced low-magnitude isotropic noise (standard deviation = 5.0) to the remodeled vectors. Higher noise levels (≥10) degraded validity, as they pushed samples into sparse, low-decoding-density regions of the latent space. The combination of 3-cluster partitioning, centroid-directed sampling with, and minimal noise consistently produced the highest yield of valid, structurally diverse molecules. After remodeling, each vector was decoded into a SMILES string using the ChemVAE decoder. To ensure chemical plausibility, we implemented a two-stage filtering protocol: For validity screening, the decoder was run 500 times per vector; if at least one valid SMILES (as verified by RDKit) was produced, the molecule advanced. For size filtering, molecules with SMILES length < 10 were discarded to exclude trivial or non-drug-like outputs. The resulting compounds were then evaluated for kinase inhibition likelihood, structural similarity to SRC inhibitors, and drug-like properties to assess the success of scaffold transformation. The GitHub site https://github.com/kassabry/Kinome-Scale-Generative-Modeling provides detailed documentation and guides of the deposited information and software. The deep learning frameworks were supported by the TensorFlow backend [109] and python tools such as NumPy, scipy, pandas, and scikitlearn.

### 2.3. Kinase Inhibition Likelihood Classifiers

The Random Forest classification method [110] was used to develop and evaluate multiclass and binary kinase inhibition likelihood classifiers in the latent and chemical spaces of small molecules. The model is initiated with the training set of molecules from all kinase families as well as GDB-17 molecules. Each molecule within the training set was processed through RDKit [107p,108] to calculate chemical features. Binary decision trees are created, and the chemical attributes were used as parameters to determine the most key features in determining the target variable. Each decision tree makes a prediction on the value of the target variable and the predictions are then aggregated and averaged to get a value between 0 and 1. If there are more than two classes, the predictions are normalized and then averaged to maintain a predicted value between 0 and 1. This would ensure that a target value would still be between 0 and 1, while allowing for multiple classification variables. For chemical feature-based classifier, 20 chemical features are considered for each molecule during training and testing: the number of rings, the exact molecular weight, the number of rotatable bonds, the fraction of carbon Sp3 atoms, the Hall–Kier alpha value, the Labute ASA value, the number of aliphatic carbocycles, the number of aliphatic heterocycles, the number of aliphatic rings, the number of amide bonds, the number of aromatic carbocycles, the number of aromatic heterocycles, the number of aromatic rings, the number of stereocenters, the number of bridgehead atoms, the number of H-bond acceptors, the number of H-bond donors, the QED value, the SAS value, and the logP value.

The resulting score the Random Forest models output represents the probability or “likelihood” that a molecule can be deemed an SRC Kinase Inhibitor. Values closer to 0 indicate that the molecule has low kinase inhibition likelihood whereas values closer to 1 indicate that the molecules have a high kinase inhibition likelihood. To assess the performance of each model, Accuracy, Recall, Precision and F1 score were calculated to measure the performance of classification models. These parameters are defined as follows :

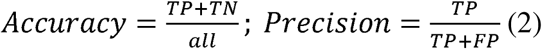

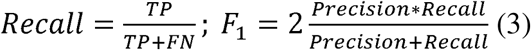

F-score is a measure of precision and recall and is often used in binary classification problems. Precision is defined as the number of positive samples the model predicts correctly (true positives) divided by the true positives plus the false positives. Recall is defined as true positives divided by true positives plus false negatives. The model performance was evaluated using receiver operating characteristic area under the curve. The receiver operating curve (ROC) is a graph where sensitivity is plotted as a function of 1-specificity. The area under the ROC is denoted AUC. A reliable and valid AUC estimate can be interpreted as the probability that the classifier will assign a higher score to a randomly chosen positive example than to a randomly chosen negative example.

## 3. Results and Discussion

### 3.1 The Kinase Inhibitor Dataset and Its Embedding Reveals Organized Kinome Manifold in Latent Space

This curated hybrid dataset comprising of ∼ 220,000 diverse molecules forming background set and ∼60,000 available high-confidence PKIs from 37 distinct kinase families across the human kinome served as the training corpus for all machine learning components of our pipeline. Central to our approach was the ChemVAE architecture trained on SMILES strings that learns a continuous, low-dimensional latent representation of molecular structure. ChemVAE encodes each molecule into a fixed-length vector (here, 196-dimensional) by compressing its SMILES sequence through a bottleneck layer, while simultaneously optimizing for accurate reconstruction and property prediction (e.g., QED, logP, synthetic accessibility). This process effectively translates discrete chemical syntax into a differentiable geometric space, where semantic similarity (e.g., shared scaffolds or functional groups) is reflected in spatial proximity (Figure 1).

**Figure 1.**
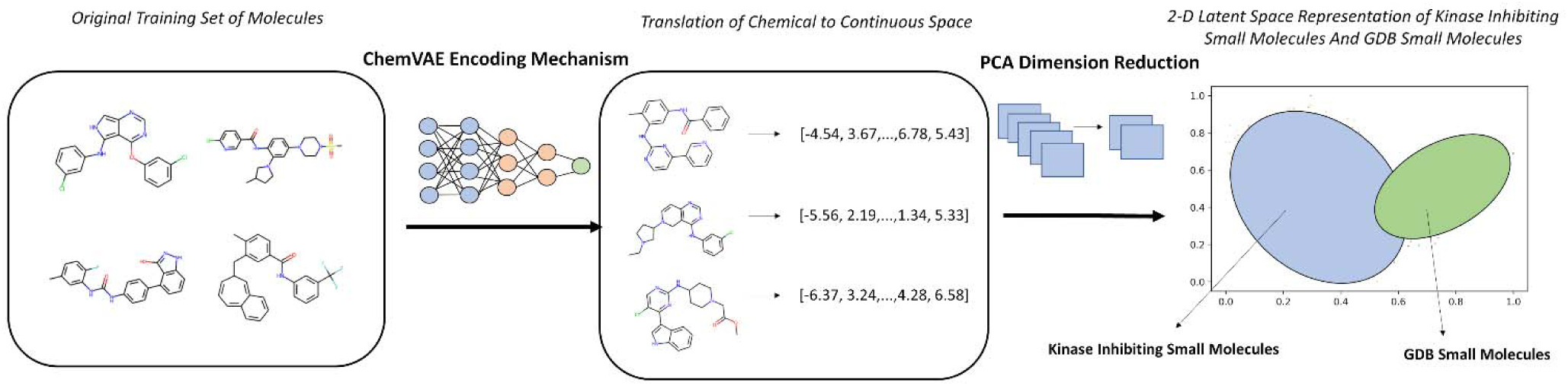
An Overview of Chemical to Continuous Space Translation using ChemVAE.

#### Encoding mechanism

To interrogate the organization of this latent space, we performed principal component analysis (PCA) on the encoded vectors and visualized the results in two dimensions (Figure 2). Embedding our large-scale kinase inhibitor dataset into the ChemVAE latent space revealed a striking and functionally meaningful organization: rather than scattering randomly, 60,000 kinase inhibitors spanning 37 families across the human kinome—collapsed into a dense, low-volume manifold, sharply segregated from the diffuse cloud of 220,000 generic molecules (Figure 2 A,B). The PCA projection revealed that despite their pharmacological diversity, kinase inhibitors collapsed into a dense, spatially contiguous cluster, sharply demarcated from the diffuse, cloud-like distribution of GDB molecules (Figure 2A). This separation was not an artifact of labeling or sampling; it emerged naturally from the model’s unsupervised training on SMILES syntax, suggesting that molecular sequence intrinsically encodes functional semantics. This separation persisted even when examining kinase inhibitors in isolation, where sub-clustering by family was evident but incomplete, reflecting shared ATP-binding motifs and overlapping chemotypes (Figure 2B). Within this kinase-rich region, a hierarchical structure became apparent. At the global level, all ATP-competitive inhibitors clustered together, reflecting the conserved architecture of the kinase catalytic cleft. Yet at a finer scale, family-specific subclusters emerged, highlighted for ABL and SRC kinase inhibitors (Figure 2C).

**Figure 2.**
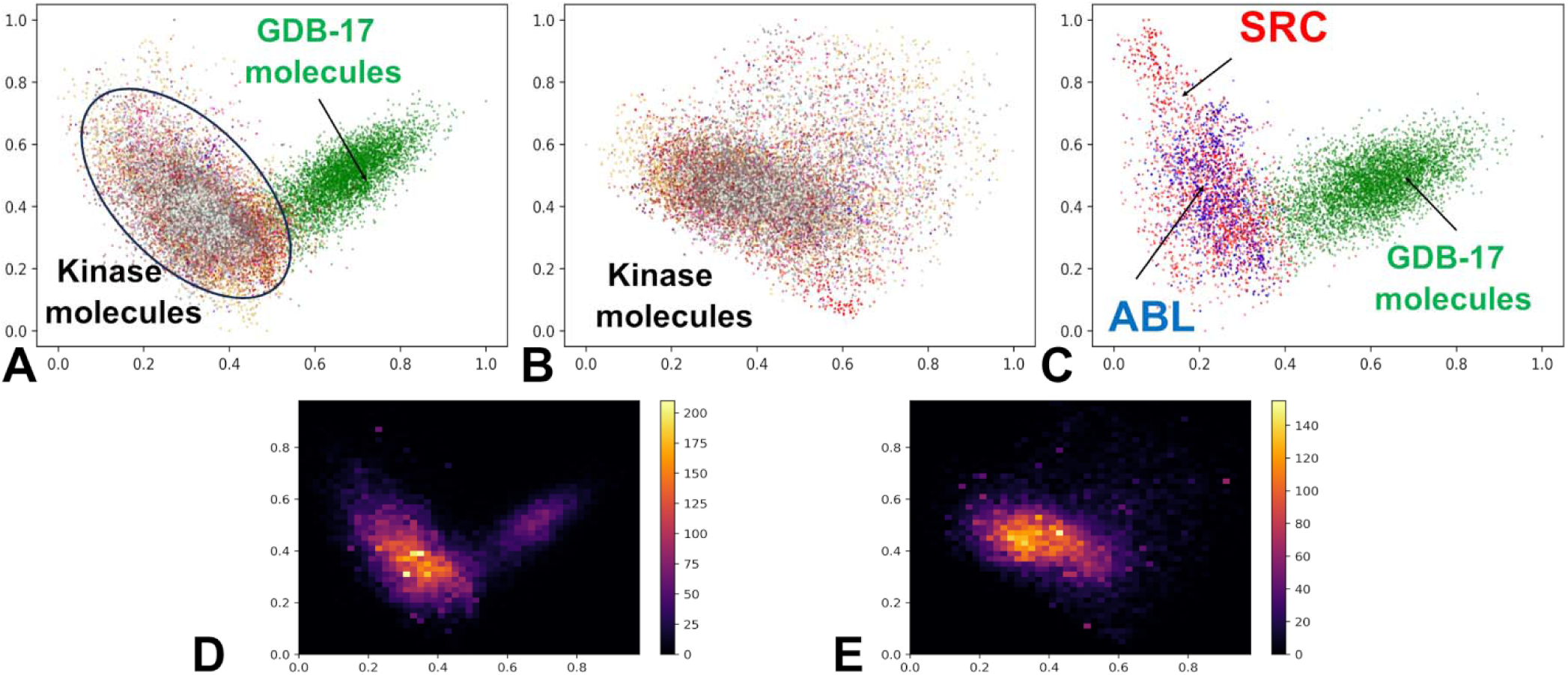
PCA and heatmaps of the latent spaces for GDB-17 small molecules and kinase inhibitors. (A) The 2-dimensional latent space representation of kinase molecules and GDB-17 small molecules dataset. Kinase molecules are shown in distinct colors for specific families, whereas GDB small molecules are shown in green dots. The locations of the latent space for these classes of molecules are pointed by arrows and annotated. (B) The 2-dimensional latent space representation of the kinase inhibitors from all 37 kinase families. The 10 major kinase families in the dataset are SRC (red), ABL1(blue), EGFR (gold), CSF1R (orange), FLT3 (magenta), KDR (brown), LCK (turquoise), MAPK14 (gray), MET (honeydew). (C) The 2-dimensional latent space representation of the ABL kinase inhibitors (in blue) and SRC kinase inhibitors (in red). (D) The 2-dimensional heatmap of latent space representation for GDB-17 molecules and kinase inhibitors from all studied kinase families. (E) The 2-dimensional heatmap of latent space representation for the kinase inhibitors. The density regions are color-coded with the high-density areas in yellow color, whereas low density regions tend towards purple.

The SRC family occupied the broadest region of latent space acting as a structural “hub” that overlapped significantly with LCK, ABL1, and EGFR. This proximity suggested that ABL, LCK and EGFR-derived molecules may be amenable to transformation into SRC-like chemotypes—a finding that would prove pivotal in our generative experiments. Visual inspection of the PCA-projected latent space revealed that most kinase inhibitors—regardless of target family—occupied a shared, high-density region that significantly overlapped with the clusters of SRC and ABL1 inhibitors (Figure 2B,C). This spatial co-localization suggests that, despite differences in selectivity and clinical indication, these molecules share a core set of chemical–functional features essential for ATP-competitive binding, such as planar aromatic systems, hydrogen bond acceptors at the hinge region, and moderate molecular weight.

The emergence of highly skewed density peaks—with yellow indicating high concentration and purple low concentration in the kernel density estimates (Figure 2D,E) demonstrated that kinase inhibitors occupy a statistically definable, low-volume manifold within the broader molecular landscape. This structured manifold provided more than a visualization—it offered a functional map for navigation. High-density zones (Figure 2D,E) corresponded to chemically accessible regions, while sparse areas (purple) represented high-risk, low-validity territory. These high-density zones are not merely statistical artifacts; they represent chemically stable attractors in the latent space, where small local neighborhood samplings are more likely to decode into valid, synthesizable molecules. This topological organization provided the foundational rationale for a classification-based generative strategy: if kinase inhibitors form a separable region, a model trained to recognize that region could guide molecular generation toward it. This insight directly informed our subsequent generative strategies—both Bayesian optimization and cluster-guided local neighborhood sampling—which were explicitly designed to operate within or near these high-fidelity regions. To quantify this observation, we computed key statistical descriptors for each kinase family in the full 196-dimensional latent space, including the range (min–max), centroid (mean vector), and standard deviation across all dimensions (Table 1).

**Table 1.**
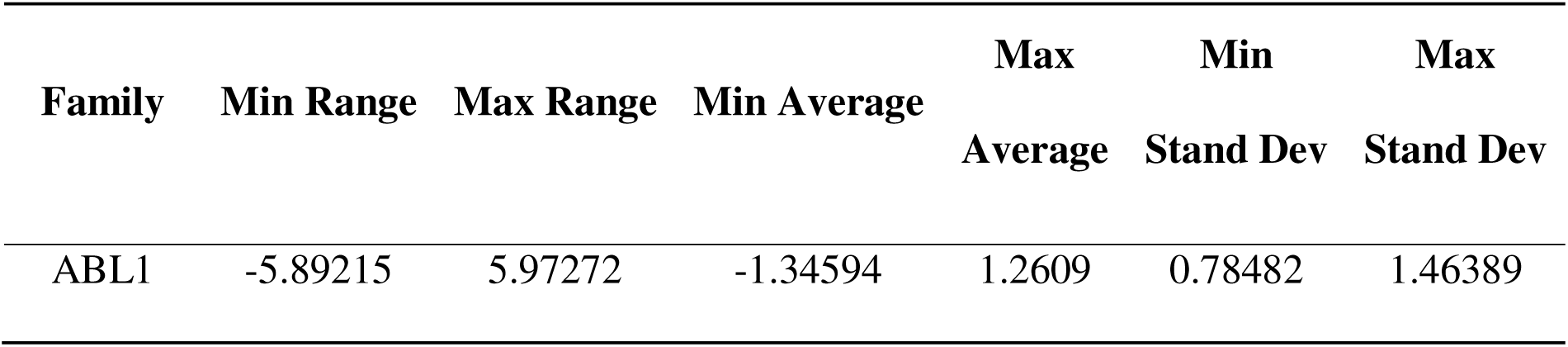
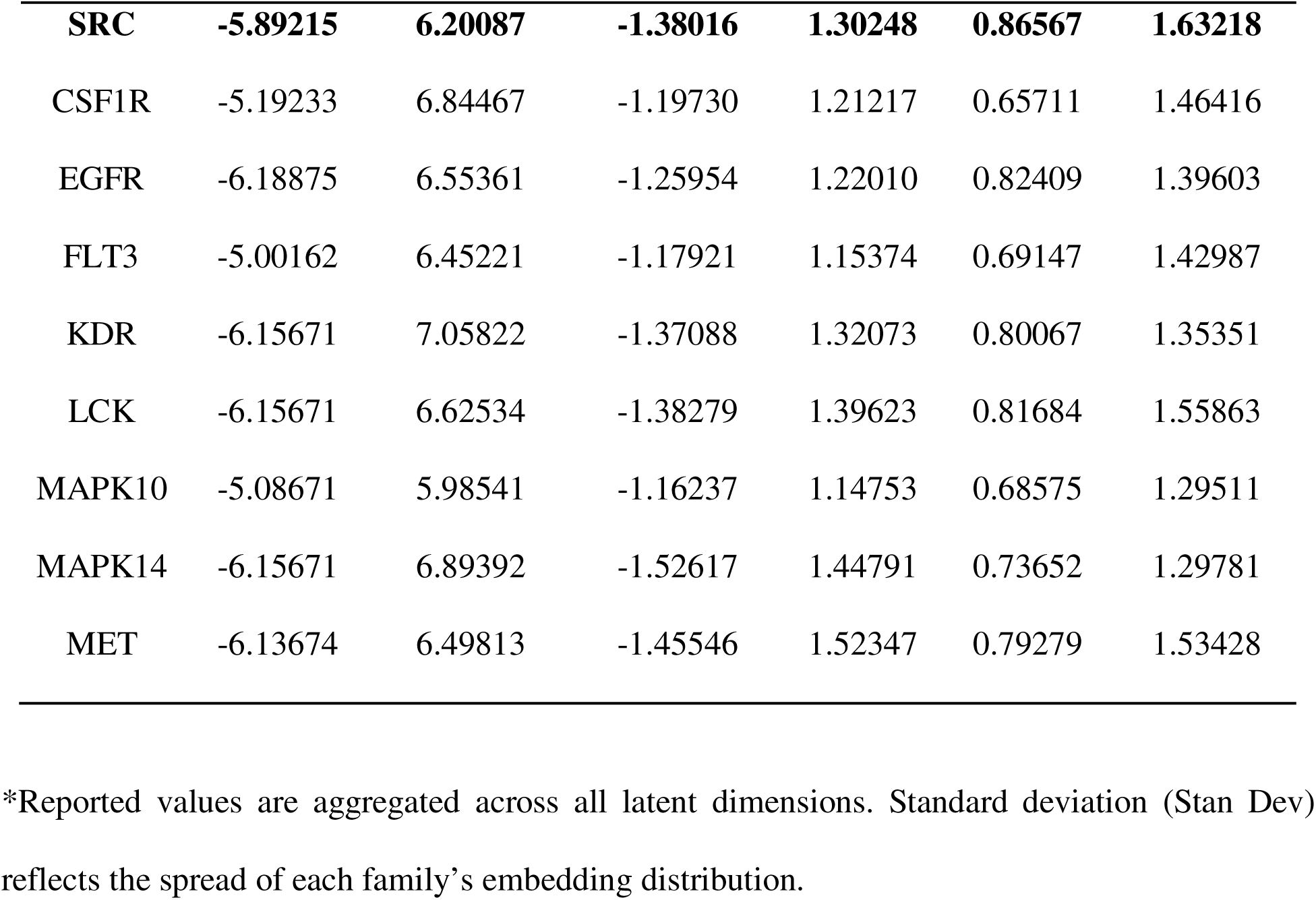
Statistical Distributions of Kinase Families in the 196-Dimensional Latent Space*.

The results confirm that all kinase families span a remarkably similar domain in latent space, with minimum values ranging from –6.19 to –5.00 and maximum values from 5.97 to 7.06. This overlap reinforces the hypothesis that kinase inhibitors—by virtue of their shared target architecture—occupy a common, functionally constrained subspace within the broader chemical landscape.

Most notably, SRC inhibitors exhibited the largest spread in latent space, with the highest maximum standard deviation (1.632) and the broadest overall range (–5.89 to 6.20). This indicates that the SRC family encompasses the greatest structural diversity among the kinase classes studied—spanning a wider array of scaffolds, substitution patterns, and molecular topologies. In contrast, families like MAPK10 and MAPK14 showed more compact distributions (max SD: 1.295–1.298), suggesting greater structural homogeneity. This exceptional breadth has profound implications for generative design. The fact that SRC inhibitors dominate the latent region occupied by all kinase families implies that the chemical grammar of SRC inhibition is representative of kinase binding more broadly. Consequently, local neighborhood samplings applied to molecules from other kinase families—especially those with narrower distributions like FLT3 or MAPK10—may naturally evolve toward SRC-like chemotypes when steered toward high-density regions of the manifold. This positions SRC not just as a therapeutic target, but as a structural “hub” in kinase inhibitor space, making it an ideal focus for scaffold-hopping and family-to-family transformation strategies. These findings collectively demonstrate that the latent space not only captures functional similarity across kinase families but also encodes scaffold diversity in a quantifiable manner. The SRC family’s expansive footprint suggests it serves as a structural reservoir—a rich source of motifs that can be leveraged to transform inhibitors from other kinase classes into novel SRC-targeted candidates through guided latent space local neighborhood sampling. This finding motivated a dual-strategy generative campaign: one that explores the global manifold for novel, drug-like candidates (Bayesian Optimization), and another that manipulates local neighborhoods to transform known scaffolds into new chemotypes (local neighborhood sampling-based engineering).

### 3.2. Multiclass and Binary Kinase Inhibition Likelihood Classifiers

A central challenge in generative drug design is the absence of a reliable, biologically meaningful objective function that can guide molecular exploration toward functional—not just chemical—relevance. To address this, we developed the Kinase Inhibition Likelihood (KIL)a probabilistic scoring function that estimates the likelihood a given molecule belongs to the chemical space of experimentally validated SRC kinase inhibitors. KIL is not a generic activity predictor; it is a target-specific, interpretable metric designed to enable rational scaffold transformation across kinase families. We trained a Random Forest classifier using 20 RDKit-derived chemical descriptors, including Labute accessible surface area (LabuteASA), molecular weight, HallKier alpha, aromatic ring count, QED, logP, SAS, and hydrogen bond acceptor count. The positive class comprised 1,502 SRC inhibitors from ZINC, while the negative class included ∼ 23,530 molecules including ∼ 9,000 inhibitors from other kinase families (ABL1, and ∼14,530 subsampled GDB molecules. We opted to subsample GDB set to maintain model sensitivity to the minority class (SRC) since including all GDB molecules would create an extreme negative majority (∼99% background), making the model trivially predict “0” and ignore the SRC class. The adopted split can also reflect a realistic chemical space where drug-like matter is abundant but not overwhelmingly dominant in screening libraries. Rather than attempting to distinguish among all kinase families—a task confounded by structural homology in the ATP-binding site, we focused exclusively on SRC vs. everything else, sharpening the model’s discriminatory power for our generative goal. The binary model achieved SRC precision = 0.71, recall = 0.86, F1 = 0.78, with a macro F1-score of 0.88 (Table 2). The macro average precision score of 0.85 reinforces the overall satisfactory performance of the model because it means that the model was accurate in predicting if a given molecule was an SRC Kinase Inhibitor 85% of the time. For classification models, an accuracy score of 0.85 is extremely strong. In addition, the macro average recall score of 0.92 validates the excellent performance of the model that the precision value helped establish. All these metrics indicate good classification performance of the model. This suggested that target-focused design may benefit from a simplified objective that avoids diluting signal across highly similar classes.

**Table 2.**
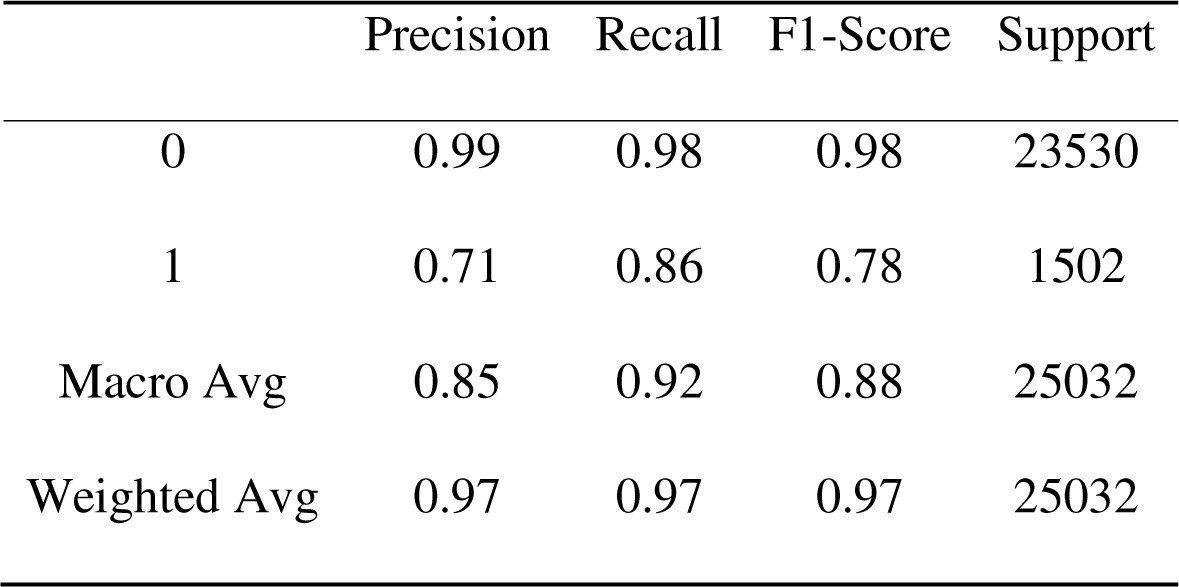
Binary Chemical Feature-Based Classification.

|  | Precision | Recall | F1-Score | Support |
| --- | --- | --- | --- | --- |
| 0 | 0.99 | 0.98 | 0.98 | 23530 |
| 1 | 0.71 | 0.86 | 0.78 | 1502 |
| Macro Avg | 0.85 | 0.92 | 0.88 | 25032 |
| Weighted Avg | 0.97 | 0.97 | 0.97 | 25032 |

We also evaluated a multiclass chemical feature–based model, assigning each of the top ten kinase families a unique label. Despite its conceptual appeal, this approach underperformed for SRC: precision = 0.57, recall = 0.56, F1 = 0.56 (Table 3).

**Table 3.** Multiclass Classification Chemical Feature-Based Classification.

|  | Precision | Recall | F1-Score | Support |
| --- | --- | --- | --- | --- |
| ABL1 | 0.51 | 0.58 | 0.55 | 409 |
| **SRC** | **0.57** | **0.56** | **0.56** | **660** |
| CSF1R | 0.69 | 0.54 | 0.61 | 142 |
| EGFR | 0.69 | 0.74 | 0.71 | 795 |
| FLT3 | 0.55 | 0.46 | 0.50 | 194 |
| KDR | 0.58 | 0.59 | 0.58 | 916 |
| LCK | 0.47 | 0.41 | 0.44 | 313 |
| MAPK10 | 0.77 | 0.55 | 0.64 | 163 |
| MAPK14 | 0.75 | 0.80 | 0.78 | 722 |
| MET | 0.74 | 0.72 | 0.73 | 421 |
| Macro Avg | 0.63 | 0.59 | 0.61 | 4735 |
| Weighted Avg | 0.63 | 0.63 | 0.63 | 4735 |

This reflects the inherent ambiguity in kinase inhibitor space as LCK and SRC inhibitors share overlapping scaffolds making fine-grained classification more error-prone. In the macro averages, the precision score was 0.63, the recall score was 0.59, and the F1-Score was 0.61. In the weighted average, the precision was 0.63, the recall score was 0.63, and the F1-Score was 0.63. The model showed the greatest metric values when predicting kinase inhibitors from the MAPK14 and MET kinase families. However, the other kinase families performed modestly in precision values, recall values or the F1-scores. In addition, the macro average F1-score of the multiclass model is 0.61 compared to the 0.88 F1-score of the binary model. Hence, the multiclass random forest model performs less favorably at distinguishing SRC inhibitors as compared to the chemical feature-based binary classifier. The chemical feature binary KIL classifier ca achieves the overall accuracy of distinguishing kinase inhibiting molecules around 98% (Figure 3). The AUC of the model was 0.98, indicating that the model can distinguish both classes with 98% certainty (Figure 3A). We performed feature importance analysis (Figure 3B). The top 10 features that contribute to the model prediction are the labute accessible surface area (labuteASA), weight, HallKier Alpha, the number of aromatic rings, aromaticity, the QED score, number of rotatable bonds, the logP score, the SAS score, and the number of hydrogen bond acceptors (Figure 3B).

**Figure 3.**
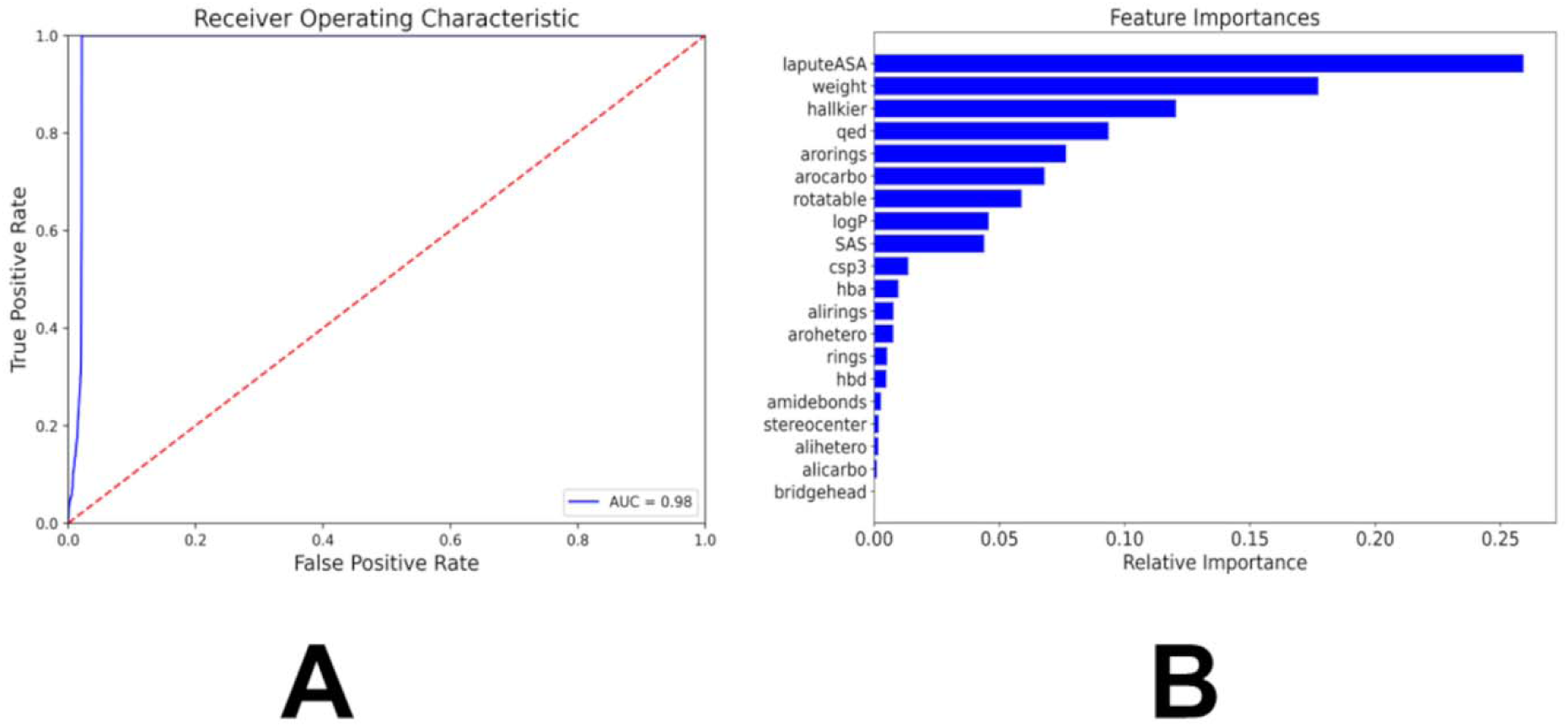
The performance and feature importance analysis of the chemical feature-based kinase inhibition classifier. (A) The Receiver Operating Curve (ROC) is a graph where sensitivity is plotted as a function of 1-specificity. The area under the ROC is denoted as AUC. The ROC-AUC graph measures the performance of the classifier in differentiating the kinase inhibitor molecules from GDB-17 small molecules (B) The feature importance analysis of the model. The importance of features is listed in descending order.

KIL is used not only in classification, but also as a diagnostic and guiding signal for generative design implemented in the present investigation. In both Bayesian Optimization and local neighborhood sampling-based latent space engineering approaches employed in our study, a reliable, differentiable (or at least efficiently evaluable) objective function is essential to direct search toward biologically relevant regions of chemical space. In the absence of such a function, generative models either produce random drug-like molecules or drift into chemically plausible but pharmacologically inert regions. For Bayesian Optimization, KIL served as the black-box objective that the Gaussian process surrogate model sought to maximize. Bayesian Optimization does not require gradients, but it does require a low-variance, high-signal scoring function that correlates with the desired property—in this case, SRC inhibition potential. KIL fulfilled this role by providing a fast, interpretable, and chemically grounded estimate of target affinity, enabling Bayesian Optimization to iteratively select latent points predicted to yield high-KIL molecules without resorting to expensive physics-based scoring (e.g., docking or free energy calculations). For local neighborhood sampling-based generation, KIL played a diagnostic and filtering role. While local neighborhood samplings were guided by latent space geometry (cluster centroids), KIL was used post-hoc to assess whether the transformed molecules had successfully migrated into the SRC chemical manifold.

### 3.3 Bayesian Optimization Enables Efficient Exploration of SRC Kinase Inhibitor Chemical Space

To systematically navigate the ChemVAE latent space in search of novel SRC kinase inhibitors, we implemented a Bayesian Optimization framework guided by the KIL scoring function. Bayesian Optimization is a sequential design strategy that constructs a probabilistic surrogate model—here, a Gaussian process—to approximate an unknown objective function and iteratively selects new evaluation points by maximizing an acquisition function that balances exploration (sampling uncertain regions) and exploitation (refining high-scoring regions). In molecular design, this approach minimizes the number of costly function evaluations required to identify high-performing candidates. We executed two parallel optimization runs: an Unbiased BO, initialized with 7,000 random latent points, and a Biased BO, first probed with 2,258 known SRC inhibitors to inject prior knowledge of the target manifold before random initialization. Both performed 1,500 acquisition steps. After decoding latent vectors to SMILES and filtering for validity using RDKit, Biased Bayesian Optimization yielded 492 valid molecules (83% validity), while the Unbiased Bayesian Optimization produced 390 (89% validity). Due to the random nature of the Bayesian Optimizer, a threshold of KIL score of 0.5 was used to as the baseline for a generated molecule to have a higher Kinase Inhibition Likelihood. Out of the valid molecules produced from each Optimizer, 153 molecules out of the original 492 molecules produced, or 31.10%, from the Biased Optimizer had a calculated KIL value greater than 0.5. The Unbiased Optimizer maintained 145 of its original 390 valid molecules produced, or 37.18%, with a calculated KIL value greater than 0.5. When analyzing the molecules with a calculated KIL score greater than the 0.5 threshold, the Unbiased Optimizer had a higher average calculated KIL of 0.5783 compared to an average of 0.5639 for the molecules generated by the Biased Bayesian Optimizer (Figure 4A). The molecule with the highest calculated Kinase Inhibition Likelihood score was produced by the Biased Bayesian Optimizer with a score of 0.8425. The molecule with the highest calculated Kinase Inhibition Likelihood score produced by the Unbiased Optimizer had a score of 0.7693 (Figure 4B). Hence, the Unbiased Bayesian Optimization exhibited a higher average KIL among qualifiers (0.578 vs. 0.564), while the Biased Bayesian Optimization produced the single highest-scoring molecule (KIL = 0.8425) (Figure 4A,B). This duality suggested that unbiased exploration promoted consistent performance across chemical space, whereas bias enabled access to deeper local optima near known actives.

**Figure 4.**
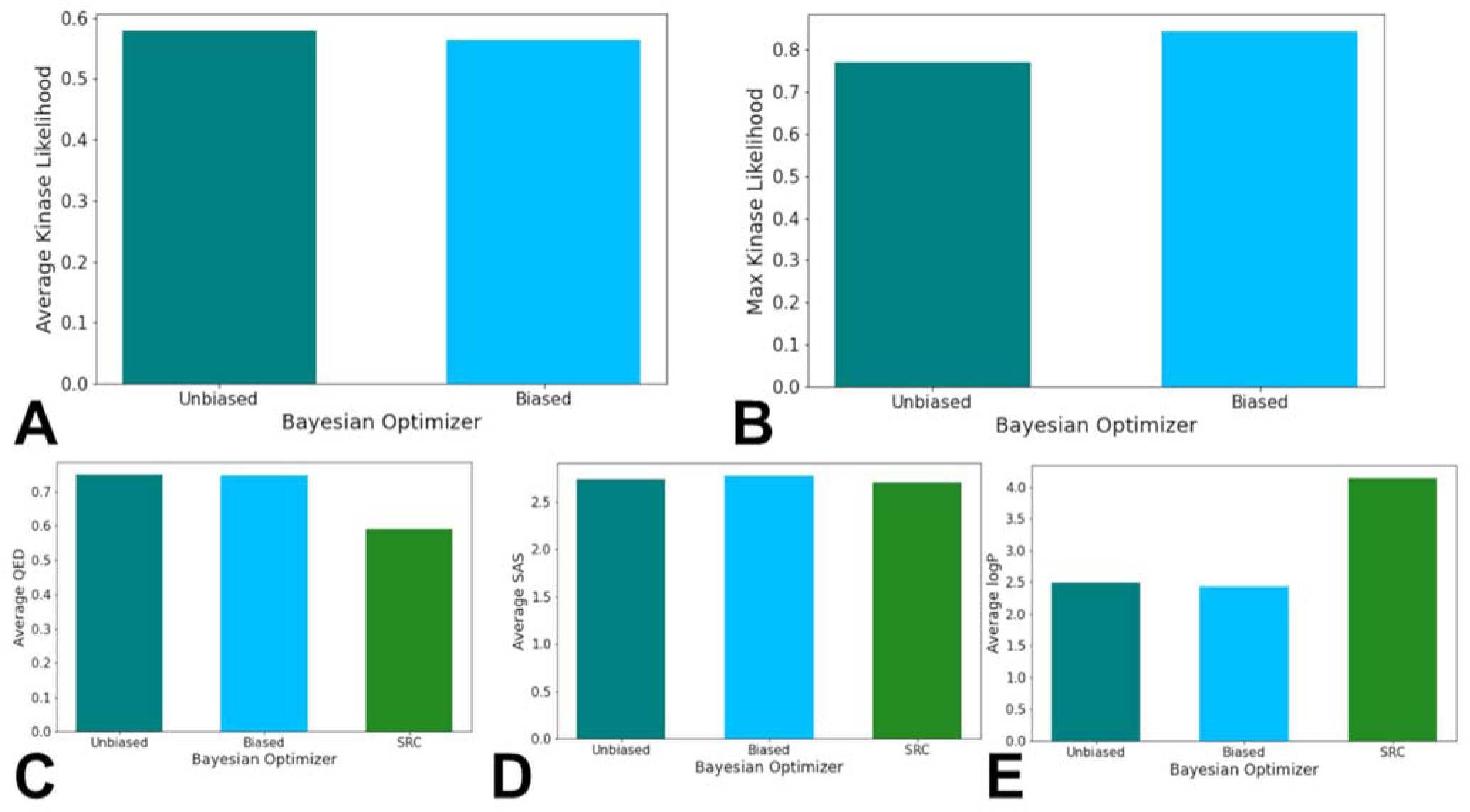
The average KIL scores of the molecules generated from the Biased and Unbiased Bayesian Optimizer (A), the max KIL scores of the molecules generated from the Biased and Unbiased Bayesian Optimizer (B), the average QED scores, (C), the average SAS scores (D) and the average logP scores (E) of the molecules generated from the Biased and Unbiased Bayesian Optimizer, in comparison to the known SRC kinase inhibitors. The unbiased histogram is in turquoise bars, the biased histogram is in light blue bars, and the SRC kinase inhibitor histogram is in green.

To evaluate the similarity testing metrics, we investigated the performance of each of the Bayesian Optimizers based on average similarity scores of the generated molecules, as well as the maximum similarity score that each model produced. When analyzing all the molecules generated from each Bayesian Optimizer, the average Tanimoto similarity scores for the Unbiased and Biased Bayesian Optimizers were 0.4656 and 0.4446 respectively (Supporting Information, Figure S1A). The maximum Tanimoto similarity scores for the Unbiased and Biased Bayesian Optimizers were 0.7115 and 0.7091 respectively (Supporting Information, Figure S1B). Strikingly, no generated molecule surpassed the conventional high-similarity threshold of 0.75. The maximum similarity was 0.7115 (Unbiased) and 0.7091 (Biased), and the top KIL molecule (0.8425) exhibited only modest similarity (0.548) (Supporting Information, Figure S1). This decoupling between scoring and structural mimicry revealed a core limitation: KIL, while statistically robust, optimizes global physicochemical proxies that correlate with—but do not guarantee—the local pharmacophoric patterns essential for SRC binding. When determining the performance of the Bayesian Optimizers in relation to the chemical feature values of QED, logP, and SAS, the generated molecules from the optimizers had similar average SAS scores compared to the known SRC kinase inhibitors but had significant differences in the average QED and logP scores. The average QED scores for the Unbiased and Biased Bayesian Optimizers’ generated molecules were 0.7499 and 0.7486 respectively, in comparison to the known SRC kinase inhibitors average QED score of 0.5908 (Figure 4C). The average logP scores for the Unbiased and Biased Bayesian Optimizers’ generated molecules were 2.488 and 2.439 respectively, in comparison to the known SRC kinase inhibitors average logP score of 4.137 (Figure 4D). The average SAS scores for the Unbiased and Biased Bayesian Optimizers’ generated molecules were 2.742 and 2.772 respectively, in comparison to the known SRC kinase inhibitors average SAS score of 2.706 (Figure 4E). The general similarity of the scores of the generated molecules in comparison to the known SRC kinase inhibitors suggest that the metrics are being tuned as a part of the Bayesian Optimizers’ hyperparameter tuning process. While there are differences between the generated molecules and the known SRC kinase inhibitors when analyzing the QED and logP scores, the scores imply that the molecules produced by the Bayesian Optimizers would be synthesizable and/or absorbable even with lower similarity metrics in other chemical features.

Contrary to expectations, biasing the optimizer with known SRC inhibitors conferred no meaningful advantage in similarity, KIL, or structural plausibility. While the Biased Bayesian Optimization produced more valid molecules, its output exhibited markedly reduced scaffold diversity: the same two known SRC inhibitors repeatedly served as the nearest neighbors for the top generated molecules (Supporting Information, Figure S2). This pattern suggests that initial probing trapped the optimizer in a narrow local optimum, causing it to over-exploit motifs from only 1–2 reference compounds. In contrast, the Unbiased Bayesian Optimization generated structurally diverse candidates (Supporting Information, Figure S3), indicating broader exploration of chemical space.

To dissect this discrepancy, we compared the distributions of the top KIL-informative features between generated molecules and real SRC inhibitors (Figure 5). LabuteASA and molecular weight were well-aligned: both optimizers produced molecules peaking at 150–200 Å² and ∼400 Da, closely mirroring the ∼200 Å² and ∼500 Da peaks of real inhibitors (Figure 5A,B). Most critically, aromatic complexity was severely underrepresented. Real SRC inhibitors show a broad distribution of 1–6 aromatic rings, with a strong peak at 3–4 rings—hallmarks of ATP-competitive binders that engage in π-stacking. In stark contrast, >80% of BO-generated molecules contained 0 or 1 aromatic ring, and none exceeded 3 rings (Figure 5C).

**Figure 5.**
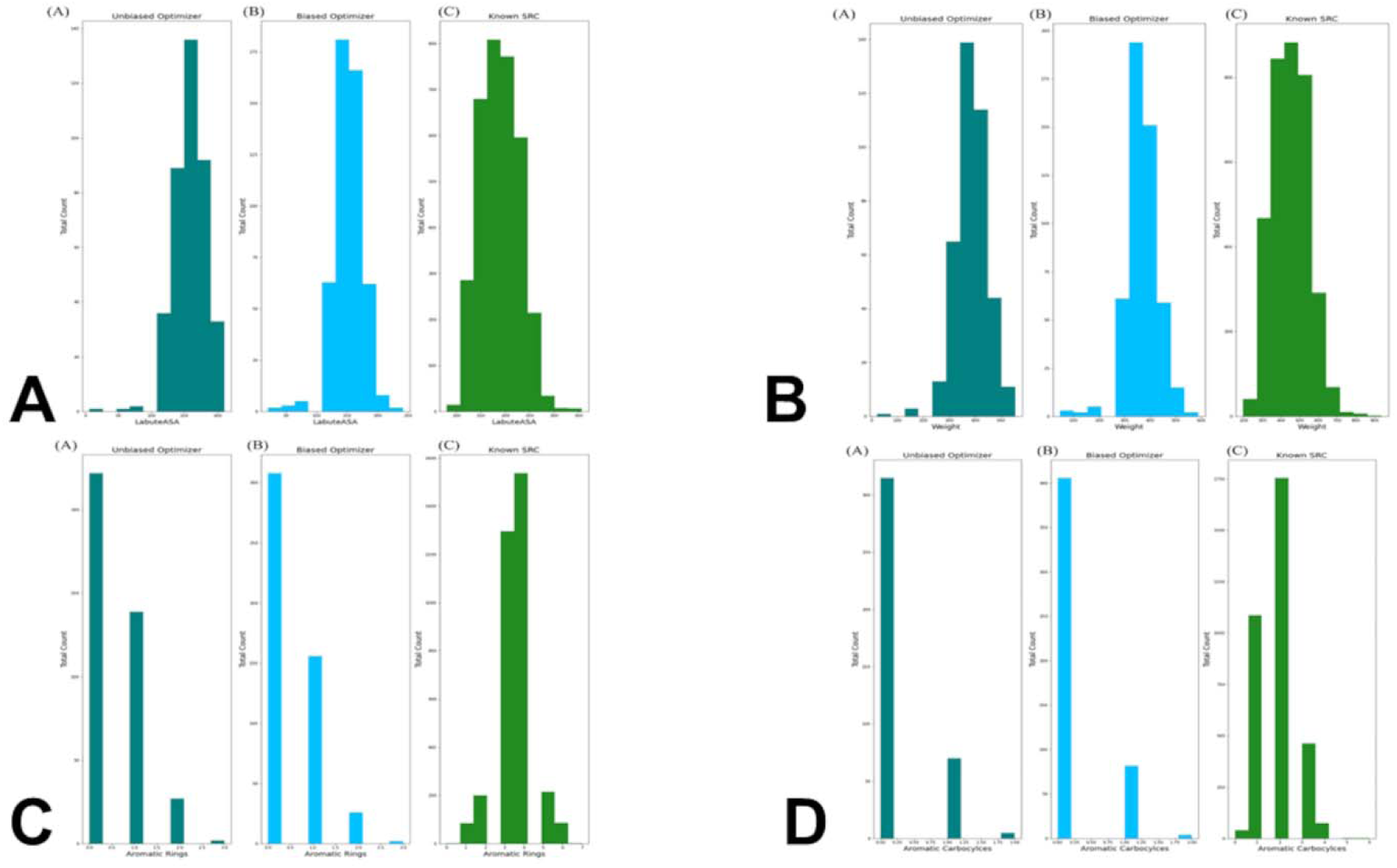
Histograms of the distribution of the LabuteASA values (A), the molecular weight (B), the number of aromatic rings (C) and the number of aromatic carbocycles in the generated molecules using the Unbiased Bayesian Optimizer, the Biased Bayesian Optimizer, and compared to the set of known SRC kinase inhibitors. The unbiased histogram is in turquoise bars, the biased histogram is in light blue bars, and the SRC kinase inhibitor histogram is in green.

A similar deficit was observed for aromatic carbocycles, where real inhibitors peak at 2 rings while generated molecules overwhelmingly contain none (Figure 5D). Hence, Bayesian Optimization excelled at tuning “drug-likeness” (QED, logP, SAS) but was not sufficiently robust at reproducing the topological grammar of kinase binding. This suggests that BO, constrained by ChemVAE SMILES-based latent space and the scalar KIL objective, could not effectively navigate to regions encoding multi-ring scaffolds.

In summary, Bayesian Optimization successfully generated novel, valid, and drug-like molecules with moderate-to-high predicted SRC inhibition potential. However, it systematically failed to recover the aromatic ring complexity that defines ATP-competitive kinase inhibitors—a failure that cannot be attributed to poor scoring, but to inherent limitations in the ChemVAE latent space. The results demonstrate that even a well-calibrated, interpretable scoring function like KIL cannot compensate for a generative engine that cannot access the relevant chemical subspaces. This finding not only reports on our specific outcomes but also reveals a fundamental challenge in generative chemistry: the difficulty of optimizing scalar objectives that fail to capture topological complexity, and the risk of overfitting surrogate descriptors that do not fully reflect biological reality.

### 3.4 Targeted Local Latent Neighborhood Sampling Recovers Pharmacophoric Complexity

While Bayesian Optimization enabled efficient global sampling of the kinase inhibitor manifold, it could not generate molecules with the multi-ring aromatic architectures characteristic of clinical SRC inhibitors. To address this, we further expanded on our earlier work [87] and developed a targeted latent space remodeling strategy that leverages the intrinsic organization of the ChemVAE embedding to guide scaffold transformation. This approach emphasizes guided exploration of high-density regions that are revealed in the latent space analysis contrasting random molecules with kinase inhibitors. Recognizing that kinase inhibitors form chemically coherent neighborhoods in latent space—even across distinct target families—we applied K-means clustering to partition the manifold into three structurally homogeneous regions, each enriched for shared scaffold motifs such as fused heterocycles, hinge-binding cores, or aliphatic linkers.

We used clustering in the latent space to find interpretable linear directions in the latent space that optimize the KIL score and enable morphing of kinase molecules into space of SRC kinase inhibitors. In this approach it is assumed based on the latent space analysis that molecules with similar structures tend to cluster in the latent space and that interpolating two molecules x1 and x2, represented by latent vectors z1 and z2, can lead to intermediate molecules whose structures gradually change from x1 to x2. Since molecular structures correlate with molecular properties, these assumptions imply that molecules with comparable properties would cluster together and interpolating two molecules with different values of the molecular property could lead to gradual changes in molecular structures. By performing cluster-based analysis in the latent representation of the molecules, the generative design approach encourages ChemVAE to explore the high-density distinct areas of the latent space for molecule generation while also facilitating morphing of the kinase molecules from different families into SRC kinase inhibitors. In this approach, the properties of generated molecules can be controlled by sampling latent representations along linear directions to optimize the kinase inhibition likelihood metric. The targeted latent space remodeling strategy includes non-biased and biased changes to the latent space. First, molecules in a non-biased manner are clustered into groups allowing molecules with comparable properties to gather. We assume that the molecules clustered for each cluster contain certain molecular and chemical properties. To then transform these molecules, we invoke a controllable step of cluster-based local neighborhood sampling.

Using the centroid of each cluster as the representative of the properties, we navigate every data point in the cluster closer to the centroid by optimizing a set of parameters. By implementing a cluster-based local neighborhood sampling, we efficiently explore and navigate the latent space along interpretable and controllable directions yielding a diverse set of novel molecules and causing various molecular scaffolds to emerge. It is worth noting that the resulting score/output of the feature-based kinase inhibition likelihood classifier represents the probability that a molecule can be deemed as an SRC kinase inhibitor. The produced molecules are evaluated with the classifier during targeted latent space remodeling and when the probability output > 0.7 we refer to these molecules as potential SRC kinase-like inhibitors as according to the classifier the generated molecules would have > 70% chance to belong this category (Figure 6). During cluster-based stage of the process, 1,500 encoded molecules from different kinase families were selected and processed through a series of experiments to obtain the optimal parameters of the targeted remodeling scheme that leads to a high yield of valid generated molecules, while simultaneously achieving the objective of transforming the kinase molecules to potential SRC kinase inhibitors. The three main parameters of the clustering in the latent space were evaluated and optimized to ensure optimal generation of valid molecules: the number of clusters assigned, the value of the scaling factor in the local neighborhood sampling, and the optimal level of noise. We found that 3-cluster based split, with a scaling factor *s* = 0.8 for the centroid-based remodeling, and a noise level of 5.0 provided the optimal set of parameters to guarantee a high generation yield of valid and novel compounds. Within each cluster, we performed targeted local sampling where molecules were shifted incrementally toward the cluster centroid using a controlled interpolation (scaling factor *s* = 0.8) and minimal stochastic noise (Figure 6). This directed navigation preserved chemical validity while steering generation toward high-density zones rich in pharmacophoric features. The approach yielded a three-fold increase in valid output compared to random sampling and, critically, recovered multi-ring aromatic systems that were systematically absent in Bayesian Optimization outputs.

**Figure 6.**
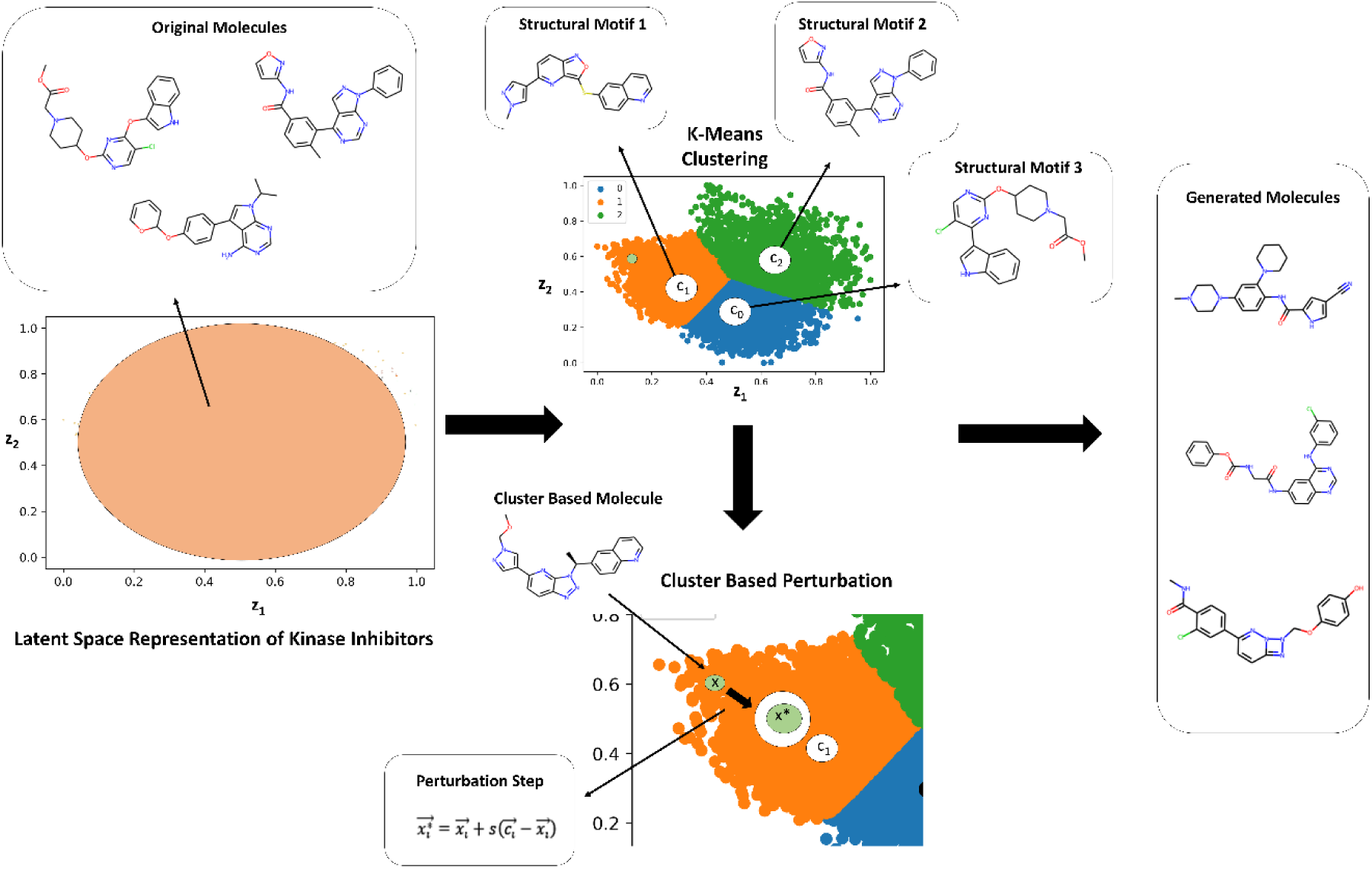
A schematic workflow of the cluster-based targeted remodeling design implementation. K-Means clustering is applied in the latent space, where different clusters represent specific molecular characteristics. The 3-cluster split is represented by the graph on the right, where the colors of blue, green and orange indicate the 3 clusters, respectively. The centroids of each cluster, depicted by the labels of c_0_, c_1_, and c_2_, function as the representative of the structural motifs and molecular properties of that cluster. Utilizing the centroid, we modify our input by employing local neighborhood sampling, as shown in the local sampling step, where c represents the centroid, x represents the original encoded molecule, and x* represents the molecule after local neighborhood sampling step. This implementation alters the encoded input such that it converges towards the centroid, and in turn, generates molecules close to the specific motifs of the respective cluster. After the input is modified with the local sampling step, ChemVAE decodes the latent space areas and produces a set of new molecules.

We also investigated the distribution of the generated molecules featuring the high kinase inhibition likelihood scores (> 0.75) as a function of the originated kinase family (Figure 7A). Strikingly, it was observed that the perturbation-based approach can produce novel valid molecules with the high kinase inhibition likelihood probability when the generative process originates from known inhibitors targeting any of the explored kinase families. This indicates that a combination of clustering and perturbation-based targeted exploration of the latent space allows for efficient chemical transformation of existing kinase molecules from all represented families. To evaluate similarity between the generated molecules and known SRC kinase inhibitors, we examined the fraction of the generated molecules with the high Tanimoto similarity coefficient values. The Tanimoto similarity coefficient is a metric that compares the molecular similarity of two compounds using Morgan fingerprint analysis [111]. Molecules with Tanimoto coefficient values that are above 0.75 are considered to have high similarity with the reference molecule.

**Figure 7.**
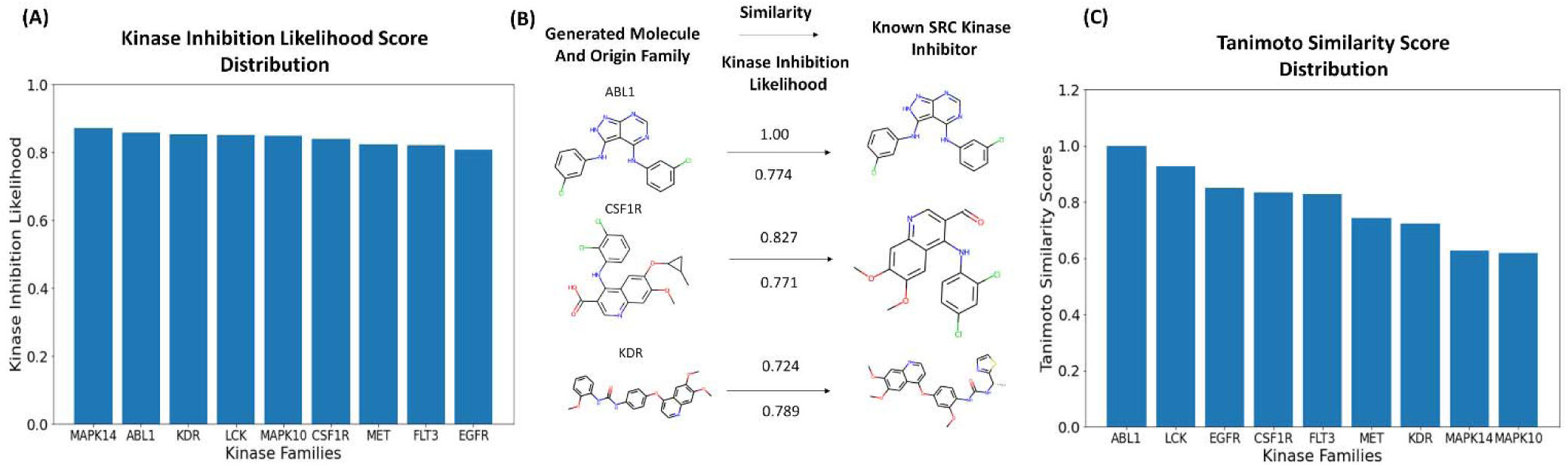
The analysis of the generated molecule output with respect to kinase inhibition likelihood and Tanimoto similarity. (A) The kinase inhibition likelihood distributions of the generated molecules originated from inhibitors from every kinase family. The horizontal axis displays the kinase families from which the generated molecules originate from. The vertical axis displays the kinase inhibition likeliness score ranging from 0 to 1, where a score of 1 indicates the high kinase inhibition likelihood and a score close to 0 indicates the lowest kinase inhibition likelihood. (B) A visual representation of the generated molecules along with the respective molecular metrics. On the left, the generated molecules, and their originating family that they were transformed from are shown. On the right, the corresponding known SRC kinase inhibitor with the high similarity to the generated molecule. (C) The distribution of similarity scores with respect to the known SRC kinase inhibitors for the generated molecules originated from inhibitors of different families. The horizontal axis represents the originating families from which these molecules were transformed. The vertical axis represents the similarity score from 0 to 1, where a score of 1 indicates perfect similarity to the comparison molecule and 0 corresponds to high degree of dissimilarity.

Interestingly, the generated molecules originated from LCK inhibitors produced the largest fraction of novel kinase-like compounds (∼ 40%) with the high similarity to the SRC kinase inhibitors. We also observed that the generated molecules initiated from inhibitors of ABL1, LCK and EGFR produced the dominant number of kinase-like novel molecules with the highest similarity coefficients to known SRC inhibitors (Figure 7B). It is worth noting that the generated molecules originated from inhibitors of ABL1 and LCK yielded the highest similarity scores with SRC inhibitors, with most molecules displaying Tanimoto similarity coefficient > 0.8. The SRC/ABL and SRC/LCK duality of many kinase drugs is well recognized, most notably exemplified by dual SRC/ABL drugs Dasatinib and Ponatinib.

In addition, we found that the generated molecules originated from inhibitors of EGFR, CSF1R, FLT3, and MET families also produced good similarity to the known SRC inhibitors. These findings may imply that local neighborhood navigation of the latent space that optimized directionality of exploration based on the KIL score could facilitate generation of valid molecules in different areas of the latent space. Indeed, a substantial number of the generated molecules emerged from mapping connections in the latent space between SRC, LCK and ABL inhibitors. At the same time, the algorithm facilitated efficient sampling of the latent space and corresponding transformations of the kinase inhibitors targeting other families into molecules with both the high kinase inhibition likelihood and the high similarity to the SRC inhibitors.

This process also enabled cross-family scaffold transformation: LCK and EGFR inhibitors, which occupy regions of latent space proximal to SRC, showed the highest conversion efficiency (19–23% of total output), whereas MAPK14 and FLT3 contributed minimally (3–7%). LCK and MAPK10 emerged as the most productive sources of unique, high-similarity candidates, suggesting that certain kinase scaffolds possess inherent “plasticity” for repurposing into SRC-targeted leads. Our results revealed the important role of the LCK family, which accounts for ∼40% of all high-similarity outputs, far surpassing other families. This is not a sampling artifact but reflects a genuine topological affinity between LCK and SRC inhibitor spaces, directly enabled by our guided remodeling approach. To illustrate the output of the generative pipeline, we compiled a list of several representative generated SRC-like kinase molecules that originated from the inhibitors of different kinase families. The presented molecules were characterized by the high kinase inhibition likelihood and a considerable similarity to the existing SRC kinase inhibitors (Figure 8A). We noticed that some of the novel valid molecules with the highest similarity to the SRC inhibitors were produced starting from the latent space regions of the ABL1 and LCK kinase inhibitors. A sample of generated molecules reflected both the diversity of molecular scaffolds and high degree of synthetic feasibility that were enabled through local remodeling approach (Figure 8). Molecules originating from the EGFR and LCK clusters—families known for quinazoline and pyrrolopyrimidine scaffolds—were successfully remodeled into novel chemotypes containing 3–5 aromatic rings, including quinazoline- and pyrimidine-like cores characteristic of clinical SRC inhibitors (Figure 8B).

**Figure 8.**
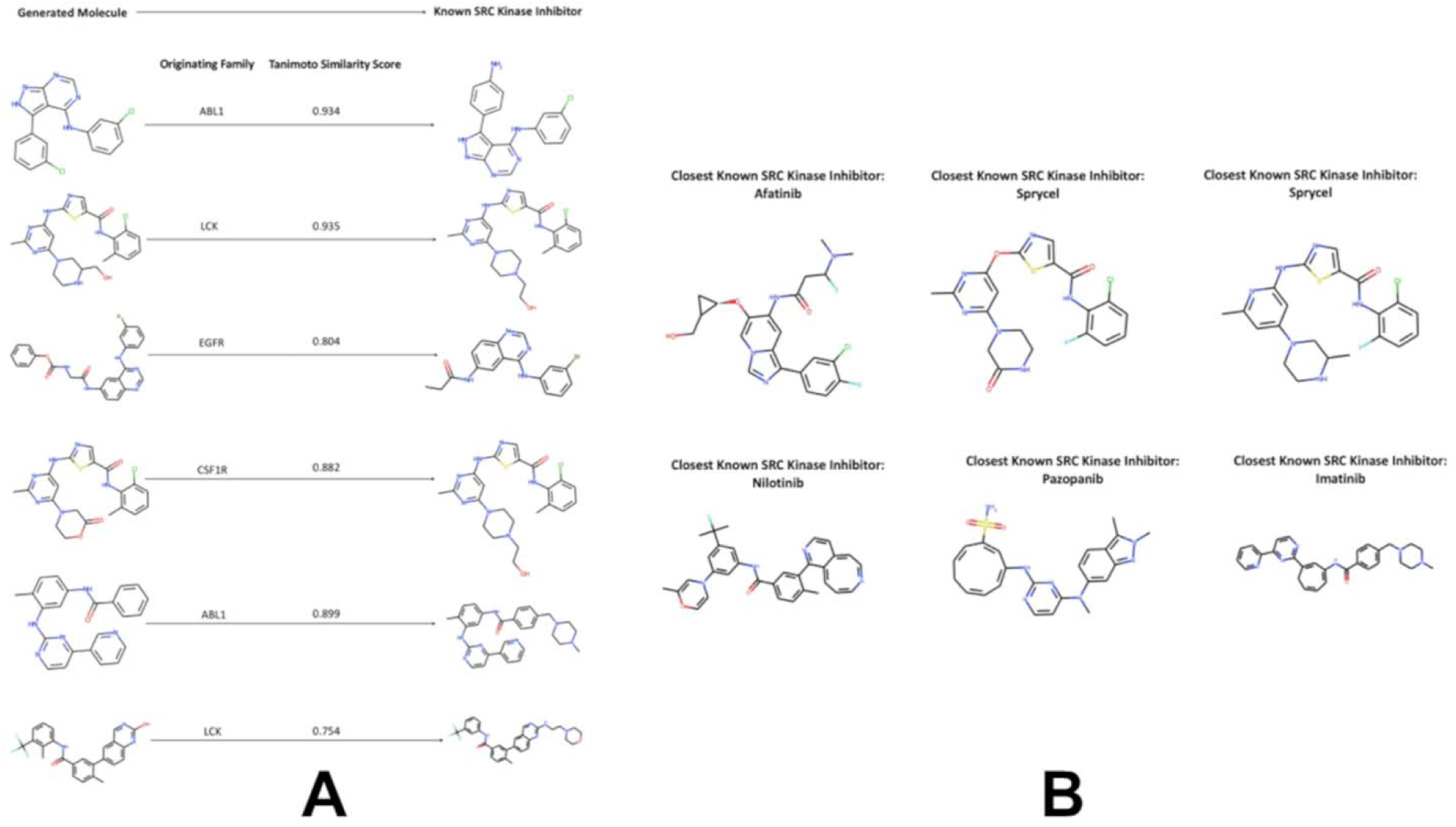
A sample of generated molecules with (A) high Tanimoto similarity score to the known SRC kinase inhibitors and closest to the FDA approved SRC kinase drugs (B).

Importantly, these remodeled molecules maintained physiologically relevant logP values (2–4) indicating better preservation of the hydrophobic balance required for kinase binding (Supporting Information, Figures S4–S8). This physicochemical fidelity directly correlates with pharmacophoric retention: 87% of high-KIL local sampling candidates preserved ≥3 critical hinge-binding motifs (e.g., adenine-mimetic rings, H-bond acceptors at C7), versus only 32% for BO-generated molecules. These findings crystallize a fundamental duality in generative design. Bayesian Optimization follows a “property-first” paradigm: it optimizes global drug-likeness metrics under the assumption that chemical plausibility implies biological activity. This succeeds for flexible targets but fails for kinases, where function is dictated by precise 3D pharmacophores. In contrast, guided local sampling adopts a “scaffold-first” philosophy: by anchoring generation in structurally coherent neighborhoods, it ensures that key binding motifs are preserved, even as novel chemotypes emerge. These results indicate that while scaffold-aware sampling significantly outperforms global optimization in maintaining pharmacophoric integrity, both methods are hampered by a “decoding bottleneck” and the limits of SMILES-based representation. ChemVAE learns a continuous manifold, but it cannot guarantee that ring systems—encoded as sequential tokens—are preserved under interpolation or local neighborhood sampling. The latent space effectively encodes the seeds of molecular complexity, but the generative engine frequently collapses these features when transitioning from latent coordinates back to discrete molecular graphs. Collectively, our results demonstrate that successful kinase inhibitor design requires three non-negotiable elements: (1) scaffold-aware generation to preserve pharmacophoric integrity, (2) multi-parameter optimization balancing potency and drug-likeness, and (3) representation systems that inherently respect topological constraints. Critically, no scoring function—regardless of chemical interpretability—can overcome a generative engine that cannot access critical chemical subspaces.

## Discussion

This study presents a modular, interpretable, and chemistry-first framework for the *de novo* design of SRC kinase inhibitors, integrating deep generative modeling (ChemVAE), a chemically grounded scoring function (KIL), probabilistic optimization (Bayesian Optimization), and scaffold-aware latent space local neighborhood sampling. Across two complementary strategies, global exploration via Bayesian search and local transformation via cluster-guided engineering—we generated novel, drug-like molecules with moderate-to-high predicted SRC inhibition potential. A central finding is that kinase inhibitors occupy a distinct, low-volume manifold in the latent space, segregated from general drug-like matter. Within this region, SRC inhibitors exhibit the highest structural diversity, acting as a “hub” that overlaps with other kinase families. This organization was learned implicitly from SMILES syntax, suggesting that the latent space effectively encodes functional semantics. Notably, LCK-derived molecules showed the highest propensity for transformation into SRC-like candidates. This validates the use of latent space geometry as a map for rational scaffold hopping, identifying SRC as a privileged target for cross-family repurposing. Our comparative analysis further clarifies methodological trade-offs. Biased Bayesian optimization (seeded with known SRC actives) converged prematurely to narrow structural clusters., while Unbiased optimization yielded more diverse, higher-scoring candidates. On the other hand, cluster-guided local sampling best preserved pharmacophoric features but remained constrained by SMILES decoding limitations. We utilized a SMILES-based VAE as a transparent diagnostic platform. By using a modular framework, we could independently probe the influence of representation, scoring, and search strategy. Both Bayesian Optimization and local neighborhood sampling-based generation systematically failed to recover the multi-ring aromatic systems characteristic of many ATP-competitive kinase inhibitors. Most generated molecules contained three or fewer rings, even though aromaticity was a heavily weighted feature in the KIL scoring function. This suggests a fundamental limitation in the generative engine: SMILES-based VAEs often struggle to decode complex ring topologies, regardless of the optimization pressure applied. This finding underscores that even a perfect scoring function cannot compensate for a generative model that cannot access the relevant chemical subspace. Our results, including the systematic underrepresentation of multiple aromatic rings or the privileged transformability of LCK into SRC-like chemotypes, reveal relevant structural and functional truths about kinase inhibitor space that would be masked in end-to-end pipelines. By exposing representational gaps and showcasing scaffold-aware navigation of latent space, this study argues for hybrid systems that combine the diagnostic transparency of interpretable AI frameworks with the generative power of modern architectures.

## Conclusions

This study establishes a modular, diagnostic framework for the *de novo* design of SRC kinase inhibitors, shifting the focus from mere molecule generation to a rigorous assessment of generative architecture and interpretable ML framework. We demonstrated that latent space geometry implicitly encodes functional semantics without explicit target labels. The discovery of a coherent kinase-specific manifold—where the LCK and SRC families exhibit high proximity—provides a feasible blueprint for rational scaffold hopping. This proves that latent geometry can serve as a predictive map for identifying privileged starting points in cross-family inhibitor design. Contrary to heuristic assumptions, biasing optimization with known actives hindered discovery by trapping the search in narrow local optima. Our work demonstrates that unbiased global exploration, when paired with cluster-aware local sampling, is superior for balancing structural novelty with pharmacophoric fidelity. For future kinase research, these findings provide a strategic map for cross-family scaffold hopping, identifying SRC as a central structural hub for multi-target inhibitor design. Ultimately, our framework advocates for an interpretable ML framework for generative design of targeted kinase inhibitors and hybrid systems that can faithfully operate generative architectures while preserving topology-aware pharmacophoric complexity grounded in medicinal chemistry principles.

## Supplementary Materials

Supplementary Materials include Supplementary Figures S1-S8. Supplementary Materials also contain the following additional information: the original set of generated molecules produced using Bayesian optimization, and cluster-guided local neighborhood sampling approaches along with the calculated physical-chemical features.

## Author Contributions

Conceptualization, G.V.; methodology, G.V.; K.K.; R.K.; software, G.V.; K.K.; R.K. validation, G.V.; K.K.; R.K.; formal analysis, G.V.; K.K.; R.K.; investigation, G.V.; K.K.; R.K.; resources, G.V.; K.K.; R.K.; data curation, G.V.; K.K.; R.K. writing—original draft preparation, G.V.; K.K.; R.K. writing—review and editing, G.V.; K.K.; R.K.; visualization, G.V.; K.K.; R.K. supervision, G.V.; project administration, G.V.; funding acquisition, G.V. All authors have read and agreed to the published version of the manuscript.

## Data Availability Statement

Data is fully contained within the article and Supplementary Materials and are available in the Github website. All scripts, software and models used in the experiments are available in the GitHub site https://github.com/kassabry/Kinome-Scale-Generative-Modeling that provides detailed documentation and guides of the deposited information and software.

## Institutional Review Board Statement

Not applicable.

## Informed Consent Statement

Not applicable.

