## Supplementary Figures S1-S8 for "From Latent Manifolds to Functional Probes: An Interpretable, Kinome-Scale Generative Machine Learning Framework for Family-Targeted Kinase Inhibitor Design"

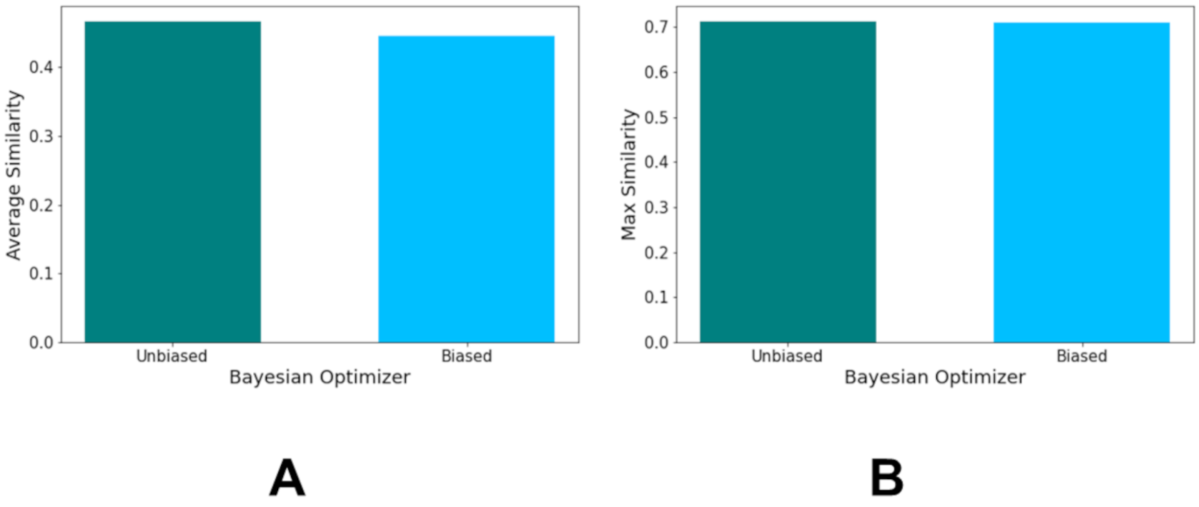


**Figure S1.** The average similarity scores to the known SRC kinase inhibitors (A) and maximum similarity to the known SRC kinase inhibitors (B) of all generated molecules from the Unbiased ( in turquoise bars ) and Biased Bayesian Optimizers ( in light blue bars)

| Generated Molecule | Known SRC Kinase Inhibitor | Tanimoto Similarity Score | Kinase Inhibition Likelihood | QED Score | logP Score | SAS Score |
|---|---|---|---|---|---|---|
| 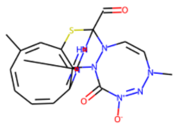 | 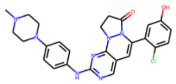 | 0.70910 | 0.57519 | 0.63193 | 1.87408 | 3.15589 |
| 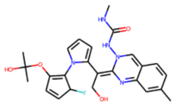 | 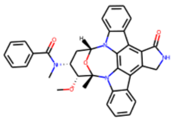 | 0.64016 | 0.75332 | 0.62254 | 4.24315 | 2.2926 |
| 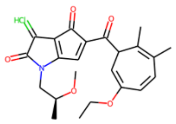 | 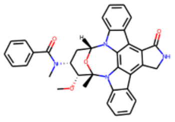 | 0.63440 | 0.58261 | 0.78757 | 0.51124 | 2.80652 |

**Figure** S2. The Top Three Molecules Generated from the Biased Bayesian Optimizer with the Closest Known SRC Kinase Inhibitors

| Generated Molecule | Known SRC Kinase Inhibitor | Tanimoto Similarity Score | Kinase Inhibition Likelihood | QED Score | logP Score | SAS Score |
|---|---|---|---|---|---|---|
| 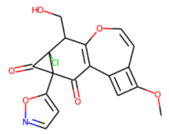 | 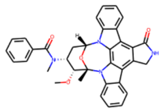 | 0.71153 | 0.58567 | 0.63763 | 3.6400 | 2.3322 |
| 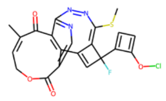 | 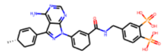 | 0.69567 | 0.59816 | 0.71623 | 4.24062 | 1.80823 |
| 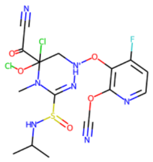 | 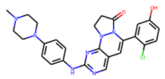 | 0.66666 | 0.60690 | 0.76593 | -0.15078 | 3.17604 |

**Figure** S3. The Top Three Molecules Generated from the Unbiased Bayesian Optimizer with the Closest Known SRC Kinase Inhibitors

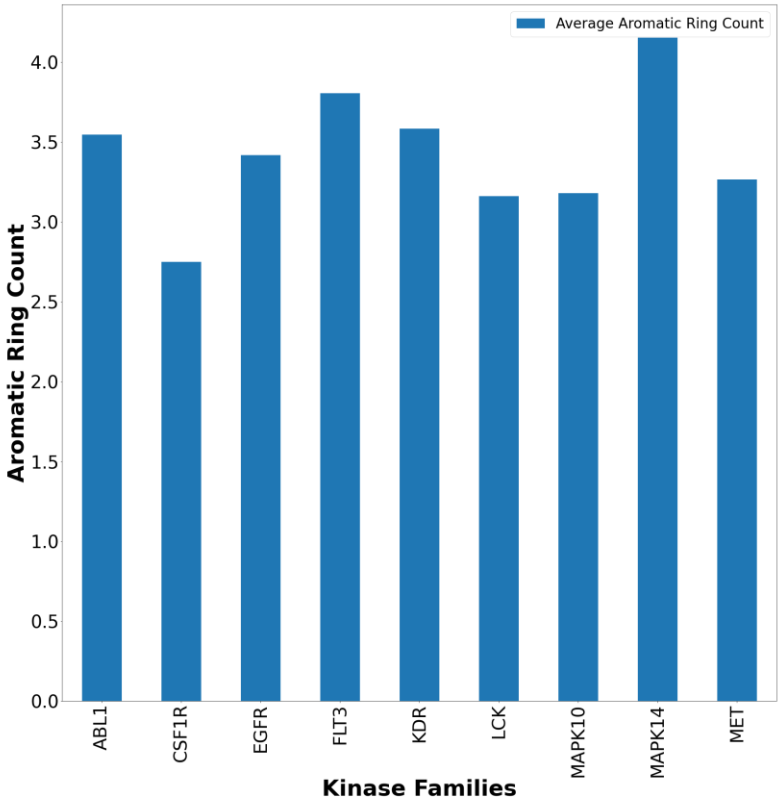


**Figure S4.** The Distribution of Average Aromatic Rings for Generated Molecules from the top 10 Originating Kinase Families.

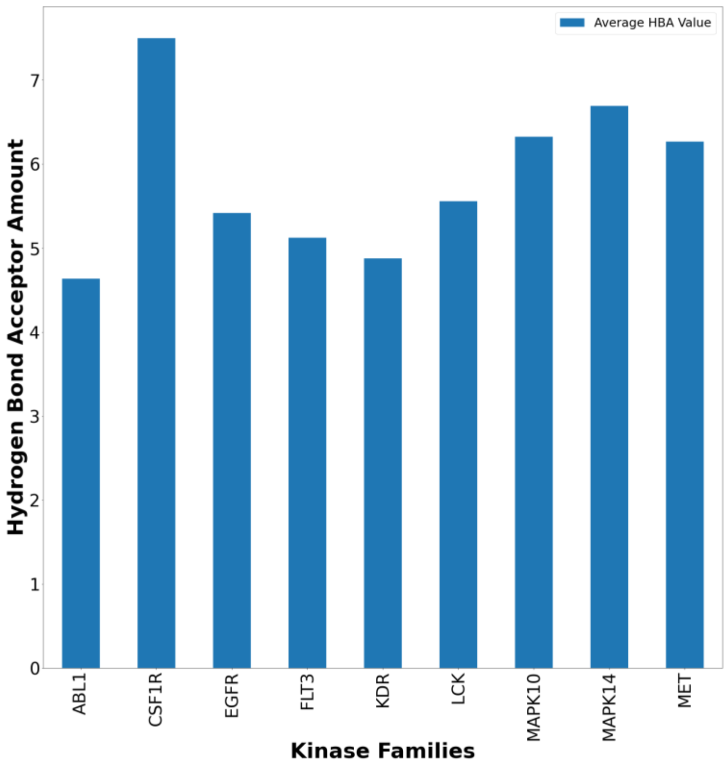


**Figure S5.** The Distribution of Average Number of Hydrogen Bond Acceptors for Generated Molecules from the top 10 Originating Kinase Families.

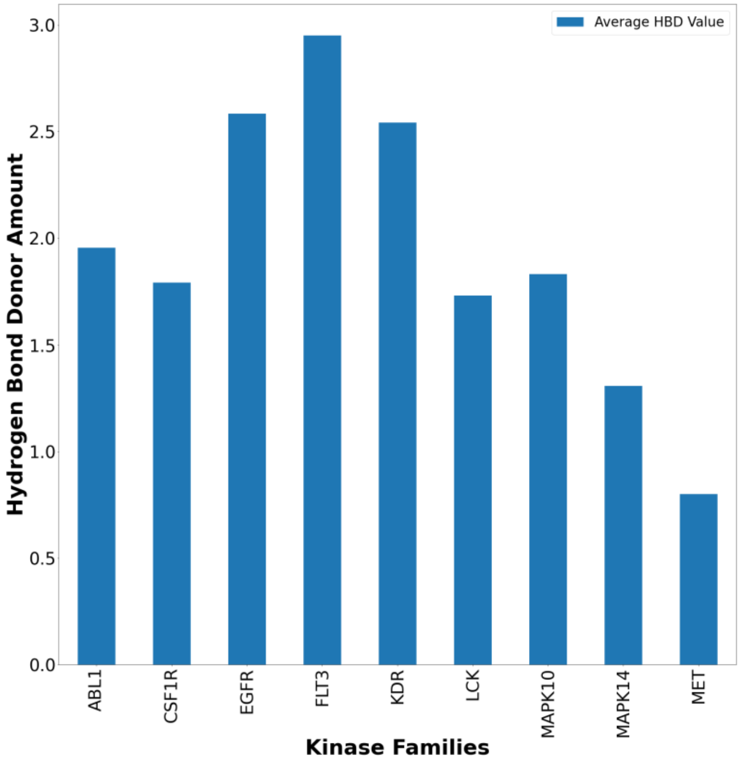


**Figure S6.** The Distribution of Average Number of Hydrogen Bond Donors for Generated Molecules from the top 10 Originating Kinase Families.

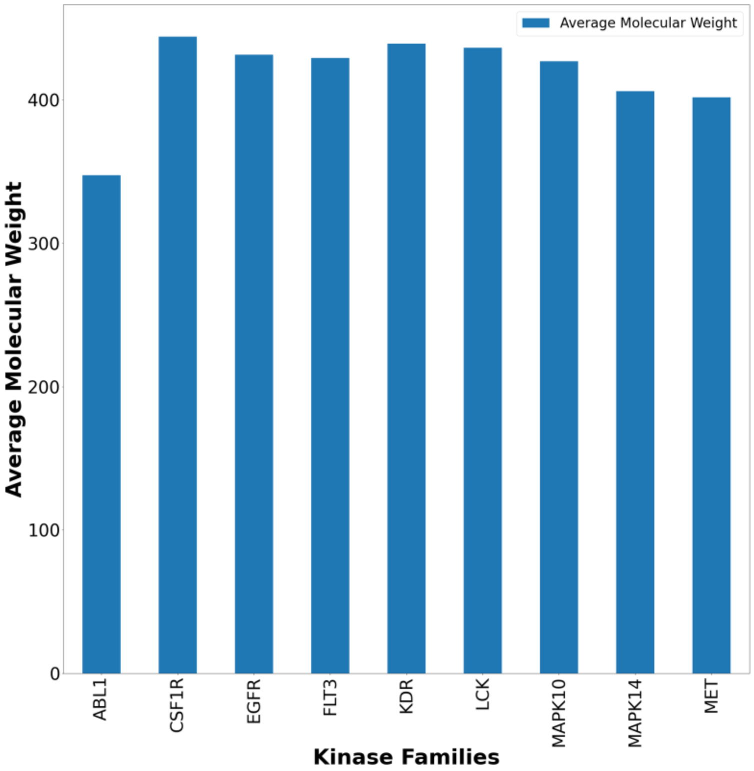


**Figure S7.** The Distribution of Average Number of Average Molecular Weight for Generated Molecules from the top 10 Originating Kinase Families.

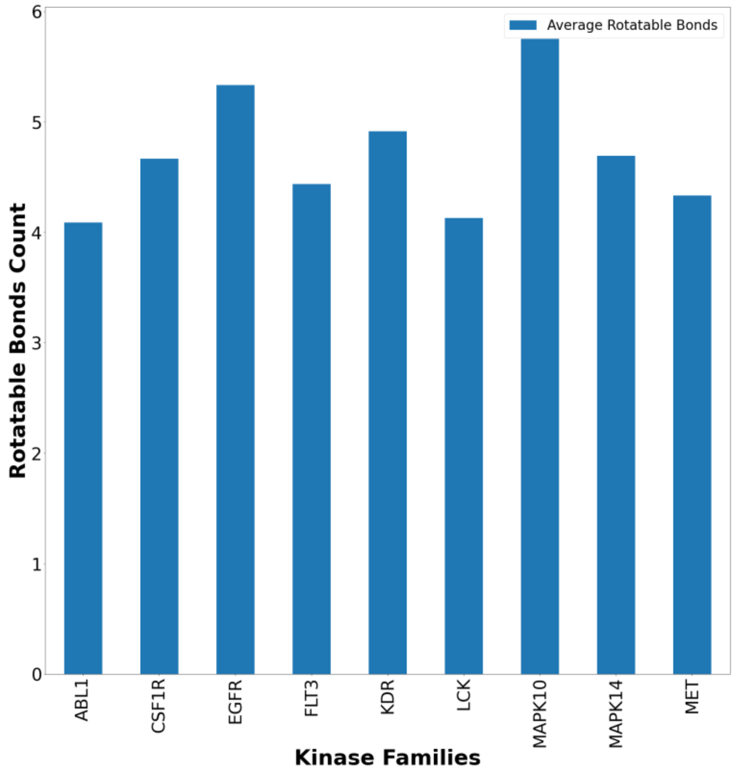


**Figure S8.** The Distribution of Average Number of Average Number of Rotatable Bonds for Generated Molecules from the top 10 Originating Kinase Families.
